# A First-in-Class, Brain-Penetrant Small Molecule Inhibitor of the NOS1AP-Tau Interaction Rescues Tau Pathology In Vivo and Improves Cognition

**DOI:** 10.64898/2026.10.01.755865

**Authors:** Ashraf N. Abdo, Sungwoo Cho, Yutong Ge, Hongliang Duan, Moustafa T. Gabr

## Abstract

NOS1AP is a genetically validated driver of tau pathology in Alzheimer’s disease (AD), yet the NOS1AP-tau interaction itself has remained pharmacologically untargeted. We report **CAPTOR-1**, a first-in-class small molecule that disrupts this interaction by binding a pocket in the N-terminal region of NOS1AP (residues 1-126). A drug-resistant NOS1AP Y16A/F42A variant that retained tau binding but exhibited a 78-fold reduction in **CAPTOR-1** affinity provides genetic evidence supporting an on-target mechanism. In human iPSC-derived neurons, **CAPTOR-1** dose-dependently reduces endogenous NOS1AP-tau association, tau seeding activity, and pathogenic tau phosphorylation, phenocopying genetic NOS1AP knockdown; these effects extend to neurons from patients with sporadic AD. Orally dosed **CAPTOR-1** crosses the blood-brain barrier and engages NOS1AP in brain dose-dependently. Chronic oral dosing rescues pathology, synaptic function, and memory in a tauopathy model (PS19) and an amyloidosis model (APP/PS1), sparing wild-type littermates in both. This work establishes pharmacological disruption of the NOS1AP-tau interaction as a viable, disease-modifying strategy for AD and positions **CAPTOR-1** as a biomarker-ready chemical probe (pTau217, neurofilament light) for translational studies.

## INTRODUCTION

Alzheimer’s disease (AD) affects tens of millions of people worldwide, and its global burden has risen sharply over the past three decades with population aging [1]. Neurofibrillary tangles composed of hyperphosphorylated, aggregated tau are a defining pathological hallmark of the disease and correlate more closely with cognitive decline than amyloid burden, making tau a central therapeutic target [2,3]. Encoded by the *MAPT* gene, tau normally stabilizes axonal microtubules; in disease, post-translational modifications and conformational changes drive its dissociation from microtubules, aggregation, and prion-like spread between neurons [2]. Tau-lowering strategies, including antisense oligonucleotides, have recently entered clinical testing; MAPT-targeting ASOs can markedly reduce CSF tau [4], and antibody-based approaches have demonstrated tau target engagement in patients [5], together providing proof that pharmacological modulation of tau biology is achievable.

NOS1AP (carboxy-terminal PDZ ligand of neuronal nitric oxide synthase, CAPON) is a neuronal adaptor protein first identified through its interaction with nNOS, where it competes with PSD-95 to redirect nNOS away from the postsynaptic density and regulate NMDA receptor-coupled nitric oxide signaling [6]. NOS1AP forms additional complexes with synapsin [7], and its interactions with nNOS occur at multiple sites, offering several potential points of pharmacological intervention [8,9]. Independent of its role in nitric oxide signaling, NOS1AP is genetically and functionally linked to a range of neuropsychiatric conditions, including schizophrenia and other psychiatric disorders [10–12], and mood, anxiety, and trauma-related disorders [13], as well as NMDA receptor-associated neuronal death [14]. Genetic and pharmacological disruption of the NOS1AP–nNOS interaction, including with the small molecule ZLc-002, produces anxiolytic effects in vivo, establishing that NOS1AP is a chemically tractable protein when approached through this interface [15].

Separately from its role at the nNOS-PSD-95 axis, NOS1AP is a direct tau-binding protein. NOS1AP accumulates in hippocampal pyramidal neurons in AD brain, and its overexpression is sufficient to drive tau hyperphosphorylation, oligomerization, and sarkosyl-insoluble tau accumulation, culminating in neurodegeneration in vivo; conversely, genetic deletion of NOS1AP ameliorates tau pathology and neurodegeneration in the P301S tauopathy mouse model [16], which was originally established by Yoshiyama et al. [17]. Mechanistically, NOS1AP promotes nitration of tau at tyrosine 29 [16], a modification enriched in neurofibrillary tangles across AD and other tauopathies [18], and associated with enhanced tau self-aggregation [19]. Nitric oxide and reactive nitrogen species more broadly have been implicated in both amyloid- and tau-driven neurotoxicity, positioning NOS1AP at a mechanistic intersection of these two pathologies [20]. Together, these findings establish the NOS1AP-tau interaction, mechanistically and structurally distinct from the NOS1AP-nNOS axis, as a genetically validated driver of tau pathology in its own right.

Despite this validation, the NOS1AP-tau interaction has remained pharmacologically untargeted. Like most adaptor proteins, NOS1AP functions through protein-protein interactions rather than a catalytic active site, and such interfaces are typically large and shallow, lacking the pockets that favor small molecule engagement [21,22]. While small molecules disrupting the NOS1AP-nNOS interaction have been developed and validated in vivo [15], no small molecule targeting the NOS1AP-tau interaction has been reported, leaving this genetically validated node pharmacologically untested.

To address this gap, we applied affinity selection-mass spectrometry (AS-MS), a label-free method for detecting direct protein-ligand interactions in solution that requires no prior structural knowledge of the binding site, well suited to adaptor proteins such as NOS1AP that lack a defined active site for conventional activity-based screening [23,24]. Our group has previously reported complementary screening platforms and hit-finding campaigns targeting the NOS1AP-nNOS interface [25–28]. Screening of diverse small molecule libraries identified a set of NOS1AP-binding hits, which we validated and ranked by microscale thermophoresis (MST). This effort yielded **NTB24**, a confirmed direct NOS1AP ligand that we subsequently optimized, through structure-activity relationship studies, into **CAPTOR-1**, a first-in-class disruptor of the NOS1AP-tau interaction. Here, we define the NOS1AP binding site engaged by **CAPTOR-1**, establish its on-target mechanism using a drug-resistant NOS1AP mutant, and demonstrate that pharmacological disruption of the NOS1AP-tau interaction reduces tau pathology and improves cognitive outcomes in human neurons and in vivo.

## RESULTS

### Discovery and optimization of a small molecule NOS1AP ligand

#### AS-MS identifies NTB24 as a direct NOS1AP ligand

NOS1AP binds tau through its N-terminal region, and genetic reduction of NOS1AP lowers tau pathology in tauopathy mice [16], yet no small molecule has been described that targets the NOS1AP-tau interaction (Fig. 1a). To identify small molecules that engage NOS1AP directly, we screened a chemically diverse library of 10,000 drug-like compounds (HitFinder collection from ThermoFisher) against recombinant NOS1AP by AS-MS, a label-free approach that requires no prior structural knowledge of the binding site (Fig. 1b). Pooled compounds were incubated with NOS1AP, protein-bound ligands were separated from free compounds by size-exclusion chromatography, and ligands were released and identified by LC-MS. The screen yielded 124 enriched compounds (1.24% hit rate), of which 50 remained after manual removal of reactive, unstable or poorly characterized molecules (Fig. 1b). Single-concentration MST validation of these 50 candidates nominated **NTB24** (Fig. 1c) on the basis of its reproducible signal, synthetic accessibility and suitability for structure-activity relationship (SAR) exploration. Concentration-dependent MST confirmed direct, saturable binding of **NTB24** to NOS1AP with a Kd of 36.7 ± 6.5 µM (Fig. 2a), within the range typical of unoptimized hits from AS-MS campaigns. Subsequent optimization of NTB24 using SAR-by-catalog approach afforded **CAPTOR-1**, with a Kd of 0.39 ± 0.17 µM (Fig. 2b).

**Figure 1.**
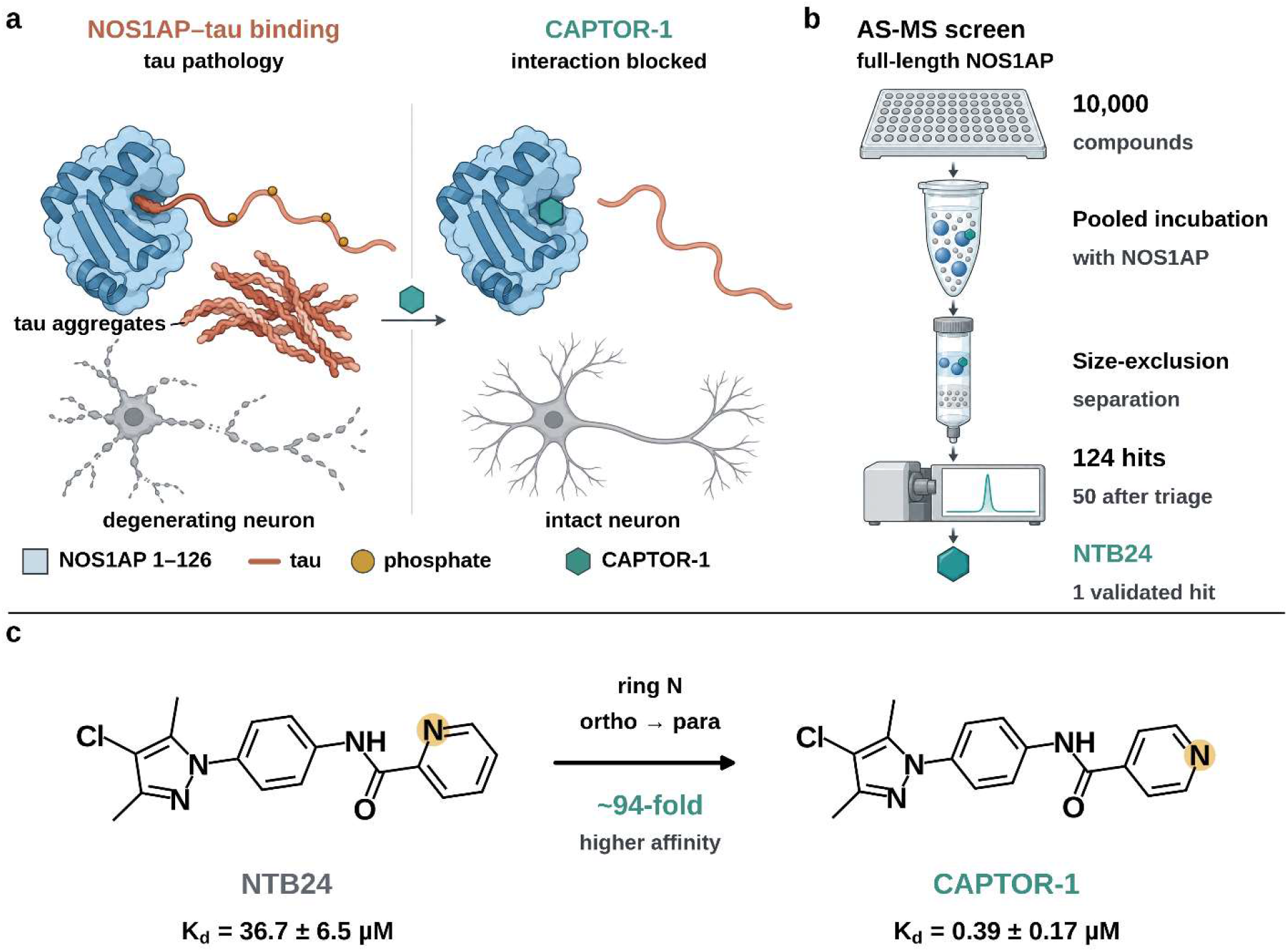
Discovery of CAPTOR-1, a small molecule disruptor of the NOS1AP-tau interaction. **a)** Schematic representation of NOS1AP (blue; N-terminal region, residues 1-126) binding the proline-rich region of tau (light orange). Binding is associated with tau phosphorylation (gold), aggregation and neurodegeneration [16]. **CAPTOR-1** (teal) occupies the NOS1AP binding site and displaces tau. **b)** AS-MS screen of 10,000 compounds against full-length NOS1AP: 124 enriched hits, 50 after manual triage, and one MST-validated hit (**NTB24**). **c)** Structures and MST Kd values (fitted value ± S.E.) of **NTB24** and its optimized analog **CAPTOR-1**, in which the pyridyl nitrogen is moved from ortho to para (∼94-fold higher affinity).

**Figure 2.**
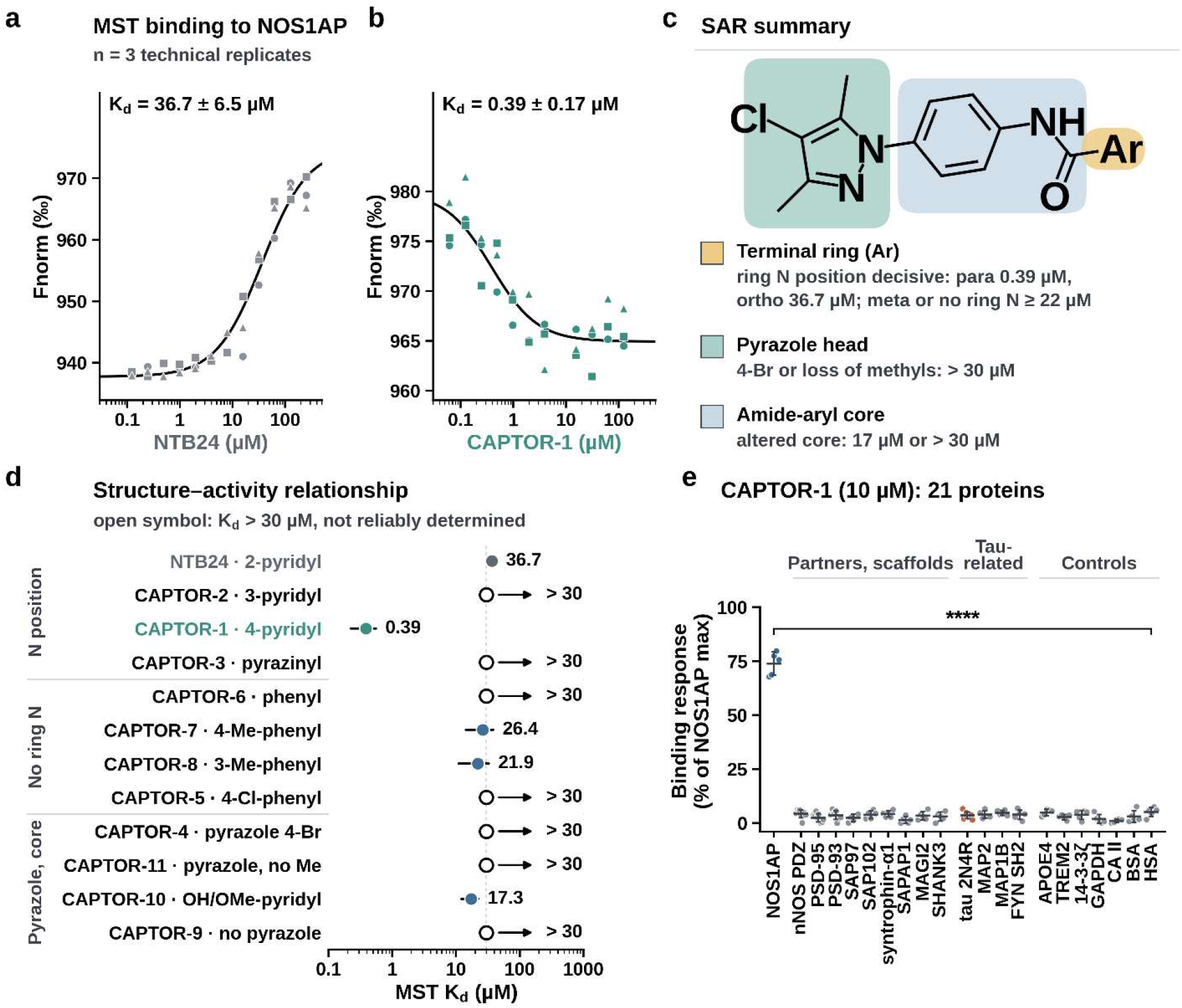
Discovery and selectivity of CAPTOR-1. **a,b)** MST dose-response of **NTB24 (a)** and **CAPTOR-1 (b)** binding to NOS1AP. All technical replicates are shown (n = 3, three symbols); lines are 1:1 binding fits; Kd is the fitted value ± S.E. **c)** General SAR summary. The scaffold is shown with the three regions varied (terminal ring, Ar; pyrazole head group; amide-aryl core) and the effect of modifications in each on MST Kd. d, MST Kd for all compounds grouped by structural change; open symbols mark curves that were unsaturated or poorly constrained (Kd > 30 µM). **e)** Binding of **CAPTOR-1** (10 µM) to NOS1AP and 20 other proteins, as % of the NOS1AP maximum response. Points are independent sensor replicates; bars are mean ± S.D. Bracket: Dunnett’s test of each protein versus NOS1AP (all *P* < 0.0001).

#### Repositioning the ring nitrogen yields CAPTOR-1

To determine the pharmacophoric requirements for NOS1AP engagement, we assembled a focused panel of eleven **NTB24** analogs varying the terminal aromatic ring, the pyrazole substituents, and the core (Fig. 2c,d; Supplementary Fig. 1 and Supplementary Table 1). The position of the ring nitrogen on the terminal pyridyl group was the dominant determinant of affinity. Relative to **NTB24**, which carries an ortho-positioned nitrogen (2-pyridyl amide; Kd = 36.7 ± 6.5 µM), relocating the nitrogen to the para position (4-pyridyl amide) yielded **CAPTOR-1**, with a Kd of 0.39 ± 0.17 µM, an approximately 94-fold improvement (Fig. 1c, 2a,b,d). The meta isomer (**CAPTOR-2**) did not produce an interpretable binding curve over the concentration range tested (Fig. 2d; Supplementary Fig. 1). Replacing the pyridyl ring with phenyl (**CAPTOR-6**) or with methyl-or chloro-substituted phenyl rings (**CAPTOR-5**, **-7**, **-8**) gave, at best, affinities in the tens of micromolar (**CAPTOR-7**, 26.4 ± 12.0 µM; **CAPTOR-8**, 21.9 ± 10.9 µM), comparable to the parent and roughly two orders of magnitude weaker than **CAPTOR-1**. Changes to the pyrazole head group or core (**CAPTOR-4**, **-9**, **-10**, **-11**) did not improve affinity, and only **CAPTOR-10** yielded a well-defined Kd (17.3 ± 5.2 µM). Curves for **CAPTOR-2**, **-3**, **-4**, **-5**, **-6**, **-9** and **-11** were unsaturated or poorly constrained and are reported as > 30 µM, not reliably determined (Fig. 2d; Supplementary Fig. 1 and Supplementary Table 1). The SAR across the three regions is summarized in Fig. 2c. A single positional change of the ring nitrogen therefore accounted for the major gain in affinity, and **CAPTOR-1** was selected as the lead for further evaluation.

#### CAPTOR-1 binds NOS1AP and none of its partners

To test whether **CAPTOR-1** engages NOS1AP selectively, we profiled it (10 µM) against 20 other proteins using biolayer interferometry (BLI): NOS1AP partners and PDZ scaffolds (nNOS PDZ domain, PSD-95, PSD-93, SAP97, SAP102, α1-syntrophin, SAPAP1, MAGI2, SHANK3), tau and tau-associated proteins (full-length 2N4R tau, MAP2, MAP1B, FYN SH2), AD-related and neuronal proteins (APOE4, TREM2, 14-3-3ζ), and unrelated or nonspecific-binding controls (GAPDH, carbonic anhydrase II, BSA, HSA) (Fig. 2e). **CAPTOR-1** gave a response of 73.9 ± 5.3% of the NOS1AP maximum with NOS1AP, versus 1.1-5.2% for each of the other 20 proteins (Dunnett *P* < 0.0001 for each; n = 5). **CAPTOR-1** did not detectably bind either of the partners whose interaction with NOS1AP is functionally relevant, nNOS or tau. For PSD-95, PSD-93, the nNOS PDZ domain and tau, responses stayed below 5% from 0.3 to 30 µM with no concentration dependence (Supplementary Fig. 2).

### CAPTOR-1 binds a defined pocket in the N-terminal region of NOS1AP

#### CAPTOR-1 binds full-length NOS1AP with submicromolar affinity

To further characterize direct binding, we measured **CAPTOR-1** binding to immobilized full-length NOS1AP by BLI. **CAPTOR-1** produced a saturable, concentration-dependent response with a Kd of 0.36 ± 0.04 µM (mean ± S.D.; Fig. 3a), in agreement with the MST-derived value (0.39 ± 0.17 µM; Fig. 2b). No response was detected on immobilized tau 2N4R or on a tag-only reference surface at concentrations up to 30 µM (Fig. 3a), consistent with the selectivity profile in Fig. 2e.

**Figure 3.**
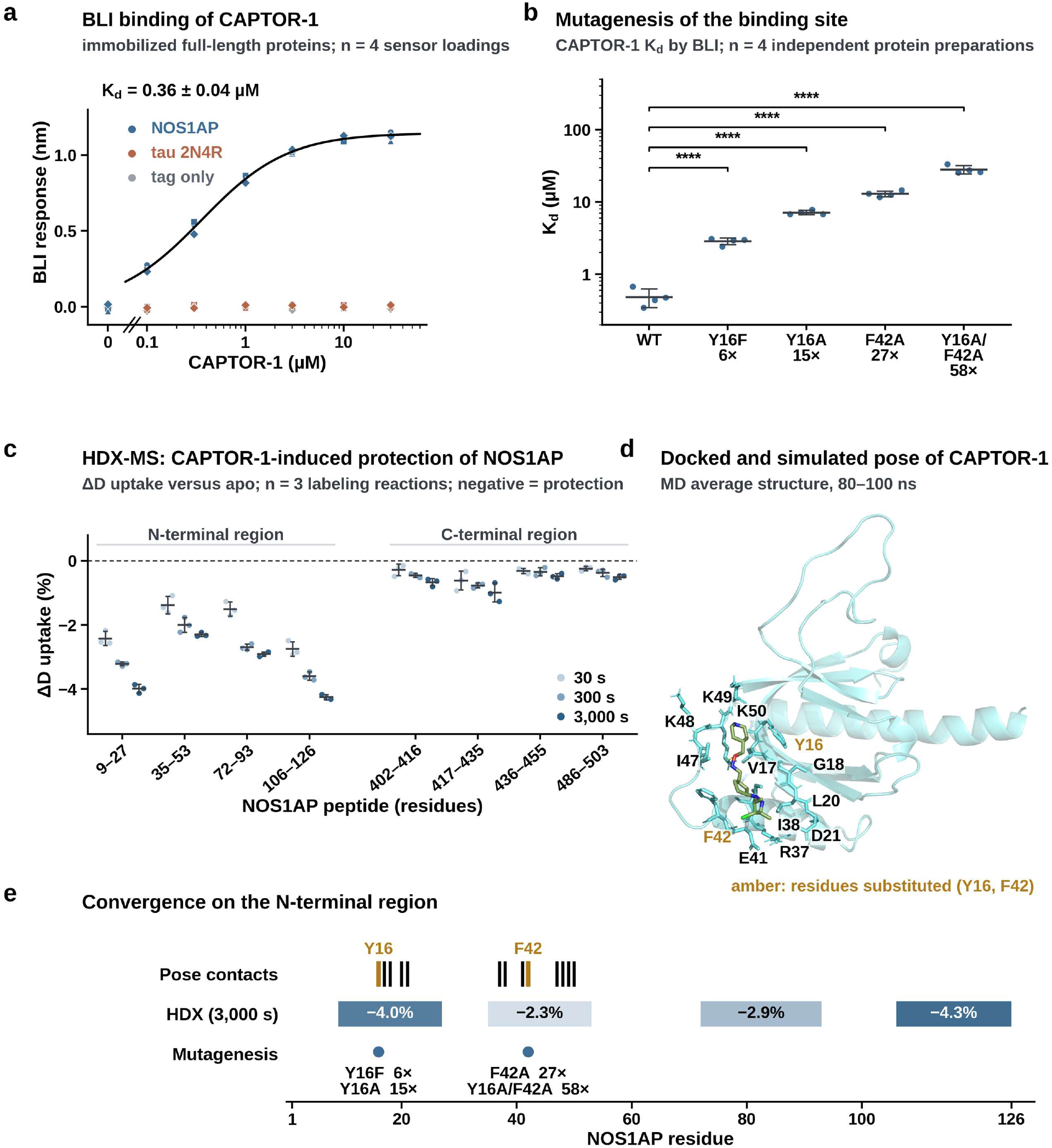
CAPTOR-1 binds a defined pocket in the N-terminal region of NOS1AP. **a)** BLI of **CAPTOR-1** binding to immobilized full-length NOS1AP, tau 2N4R and a tag-only reference. All four independent sensor loadings are shown (symbols); the line is a 1:1 saturation fit to NOS1AP; Kd is the mean ± S.D. of the four loadings. **b) CAPTOR-1** Kd (BLI) for wild-type and variant NOS1AP; points are independent protein preparations (n = 4), bars are mean ± S.D.; fold loss versus wild type is shown under each variant; brackets, Dunnett’s test on log10 Kd (**** *P* < 0.0001). **c)** HDX-MS: **CAPTOR-1**-induced change in deuterium uptake (ΔD) versus apo at 30, 300 and 3,000 s for the eight significantly responsive peptides (n = 3 independent labeling reactions; negative values indicate protection). **d) CAPTOR-1** pose in the NOS1AP homology model (MD average structure, 80-100 ns); labeled residues contact the ligand; amber marks Y16 and F42, substituted in the variants. **e)** Residue-level summary: pose contacts, HDX protection at 3,000 s (ΔD, %), and mutagenesis sites with fold loss in **CAPTOR-1** affinity.

#### Substitution of pocket residues weakens binding without disrupting the protein

To test the pocket used for docking **CAPTOR-1** to NOS1AP (see below for full details regarding the docking), we substituted two residues that the docked pose places in contact with the ligand, Y16 and F42, in recombinant full-length NOS1AP. The substitutions progressively weakened **CAPTOR-1** binding: Kd rose from 0.36 µM (wild type) to 2.9 µM (Y16F), 7.2 µM (Y16A), 13.0 µM (F42A) and 28.2 µM (Y16A/F42A), that is 8-, 20-, 36- and 78-fold losses (Dunnett *P* < 0.0001 for each; n = 4; Fig. 3b). The variants behaved normally: cellular expression, size-exclusion chromatography (SEC) monomer content and basal NOS1AP-tau interaction in cells were within 10% of wild-type values, and only F42A showed a small decrease in SEC monomer content and melting temperature (about 2 °C), which the double variant did not (Supplementary Fig. 3). The double variant Y16A/F42A therefore loses **CAPTOR-1** binding while its basal interaction with tau in cells is not significantly changed (90 ± 8% versus 93 ± 3% for wild type).

#### CAPTOR-1 protects the N-terminal region of NOS1AP from deuterium exchange

As a structural readout of the binding region, we mapped ligand-induced protection of full-length NOS1AP by hydrogen-deuterium exchange mass spectrometry (HDX-MS). **CAPTOR-1** reduced deuterium uptake in four peptides spanning the N-terminal region (residues 9-27, 35-53, 72-93 and 106-126), with protection increasing over time (−4.0%, −2.3%, −2.9% and −4.3% at 3,000 s, respectively; Fig. 3c; Supplementary Table 2). Four additional responsive peptides in the C-terminal region (residues 402-503) showed much smaller changes (−0.5% to −1.0%).

#### A docked and simulated pose of the CAPTOR-1/NOS1AP interaction is consistent with the experimental data

We docked **CAPTOR-1** into a published homology model of NOS1AP [26] with the search box centered on the predicted pocket, and refined the best pose by 100-ns explicit-solvent molecular dynamics. In the 80-100 ns average structure the ligand remains in the pocket, surrounded by residues Y16, V17, G18, L20, D21, R37, I38, E41, F42, I47, K48, K49 and K50 (Fig. 3d). Y16 and F42, the two residues substituted in the variants, are both among them. All 13 contacts lie within the first two HDX-protected peptides, whereas the protection at residues 72-126 is not explained by the pose and may reflect allosteric propagation (Fig. 3e). The parent hit **NTB24** occupies the same region and shares 8 of 13 contact residues with **CAPTOR-1** (Supplementary Fig. 4). The three systems fluctuated within a similar range over the simulation (Supplementary Fig. 5). MM/PBSA calculations favored **CAPTOR-1** over **NTB24** (−24.4 versus −14.9 kcal/mol; Supplementary Table 3), but because they come from a single trajectory per system and exclude entropy, we use them only to indicate the direction of change, not affinity. Together, BLI, mutagenesis and HDX-MS place **CAPTOR-1** in a pocket within the N-terminal region of NOS1AP, and the modeled pose is consistent with them.

### CAPTOR-1 disrupts the NOS1AP-tau interaction

#### NOS1AP binds tau through its N-terminal region and tau’s proline-rich domain

Purified NOS1AP bound tau with a Kd of 0.28 ± 0.02 µM by surface plasmon resonance (SPR; k_on_ 8.2 × 10^4^ M^−1^ s^−1^, k_off_ 0.023 s^−1^; n = 3 independent surfaces; Fig. 4a; Supplementary Fig. 6) and 0.35 ± 0.03 µM by MST (n = 3 independent protein preparations; Fig. 4b). Mapping the interface with truncation constructs showed that full-length NOS1AP bound the proline-rich domain of tau (residues 151-243) as well as full-length tau (98.5 ± 8.2% of the full-length pair), whereas the N-terminal, microtubule-binding and C-terminal tau fragments did not (about 5% or less). On the NOS1AP side, residues 1-126 were sufficient (85 ± 5% with full-length tau; 90 ± 2% with the proline-rich domain), residues 1-150 gave full binding, and residues 76-150 gave partial binding (24-27%). Residues 1-75, 127-503 and 151-503 gave no binding (about 5% or less; Fig. 4c; Supplementary Table 4). The tau-binding region of NOS1AP is therefore the N-terminal region, the same region engaged by **CAPTOR-1** (Fig. 3).

**Figure 4.**
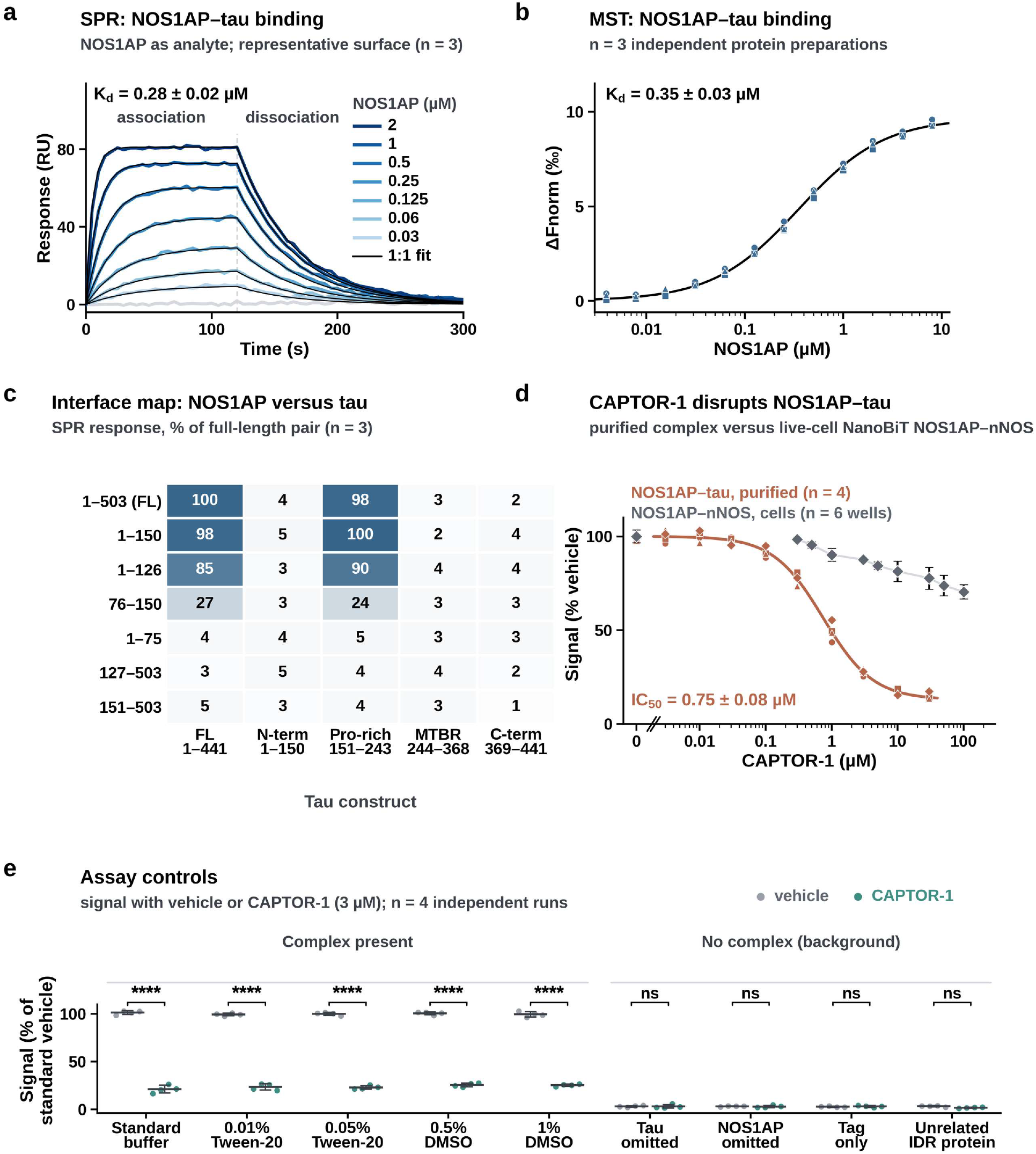
CAPTOR-1 disrupts the NOS1AP-tau interaction. **a)** SPR sensorgrams of NOS1AP (0.03-2 µM) binding to tau on a representative surface (colored lines, measured response, darker = higher concentration; light grey, buffer injection) with global 1:1 kinetic fits (black; kon and koff from the fit, Rmax fitted); the injection ends at 120 s. Kd is the mean ± S.D. of three independent surfaces (Supplementary Fig. 6). **b)** MST of NOS1AP binding to tau; all points from three independent protein preparations (symbols) with a 1:1 fit; Kd is the mean ± S.D. of the preparations. **c)** Interface map: SPR response of NOS1AP and tau constructs as % of the full-length pair (mean, n = 3). **d)** Concentration-dependent disruption by **CAPTOR-1** of the purified NOS1AP-tau complex (terracotta; all points from four independent preparations, symbols; four-parameter fit; IC_50_ is the mean ± S.D.) compared with the live-cell NanoBiT NOS1AP-nNOS signal (grey; mean ± S.D., n = 6). **e)** Assay controls: signal with vehicle or **CAPTOR-1** (3 µM) in standard buffer, with detergent or DMSO, and in background conditions without a complex (n = 4 independent runs; brackets, Welch t-tests, Holm-adjusted; **** *P* < 0.0001, ns not significant). Signals are normalized to the standard-buffer vehicle.

#### CAPTOR-1 disrupts the purified complex

**CAPTOR-1** reduced the NOS1AP-tau complex signal with an IC_50_ of 0.75 ± 0.08 µM (n = 4 independent preparations), with maximal inhibition of about 87% (Fig. 4d). At 3 µM, **CAPTOR-1** reduced the complex signal to 21-26% of vehicle in standard buffer and in the presence of 0.01-0.05% Tween-20 or 0.5-1% DMSO (Holm-adjusted *P* < 0.0001 for each). It did not significantly alter the background signal in the tau-omitted, NOS1AP-omitted, tag-only or unrelated intrinsically disordered protein conditions (Fig. 4e). The insensitivity to detergent and the unchanged background argue against colloidal aggregation or assay interference as the source of the effect.

#### CAPTOR-1 disrupts the nNOS interface only partially

Because NOS1AP also binds nNOS through a distinct interface, and small molecules directed at that interface have been reported [15,25,27], we used a live-cell NanoBiT assay of the NOS1AP-nNOS interaction [25] as a counter-screen. **CAPTOR-1** reduced the NanoBiT signal only partially, to 90.2 ± 3.5% of vehicle at 1 µM, 81.4 ± 5.4% at 10 µM and 70.4 ± 3.8% at 100 µM (n = 6; Fig. 4d; Supplementary Fig. 7). At 1 and 10 µM, the same concentrations reduced the purified NOS1AP-tau complex to 49.3 ± 4.8% and 16.7 ± 1.5% of vehicle. The parent hit **NTB24** had no effect up to 30 µM and reduced the NanoBiT signal to 62.2 ± 3.7% at 100 µM (Supplementary Fig. 7). The NanoBiT effect emerges from about 0.5 µM, in the range of the tau IC_50_. **CAPTOR-1** is therefore directed preferentially, but not exclusively, at the tau interface. The two assays differ in format (purified proteins versus live cells), so this comparison does not give an absolute selectivity ratio. Together with the absence of **CAPTOR-1** binding to the nNOS PDZ domain (Fig. 2e), the partial nNOS-interface effect is consistent with engagement of NOS1AP rather than nNOS.

### CAPTOR-1 acts through NOS1AP in human neurons

#### CAPTOR-1 reduces endogenous NOS1AP-tau association in human neurons

To test target engagement in a physiological setting, we treated human iPSC-derived neurons from three donors (two independent differentiations each) with **CAPTOR-1** and measured the endogenous NOS1AP-tau complex by immunoprecipitation and parallel reaction monitoring (IP-PRM). **CAPTOR-1** reduced the association dose-dependently, to 68 ± 9%, 51 ± 3% and 33 ± 9% of vehicle at 0.3, 1 and 3 µM (mean ± S.D. of six observations; Holm-adjusted *P* = 0.019, 0.005 and 0.018; Fig. 5a), similar to NOS1AP knockdown by CRISPR interference (CRISPRi; 28 ± 9%; *P* = 0.018). **CAPTOR-1** also reduced the endogenous NOS1AP-nNOS complex (84 ± 4%, 62 ± 8% and 48 ± 10% of vehicle; Fig. 5b). The reduction was smaller than that of the tau association at every dose in all three donors (by 16, 11 and 14 percentage points; unadjusted paired *P* = 0.036, 0.013 and 0.16), so **CAPTOR-1** engages NOS1AP at both interfaces in neurons, with a modest preference for the tau association.

**Figure 5.**
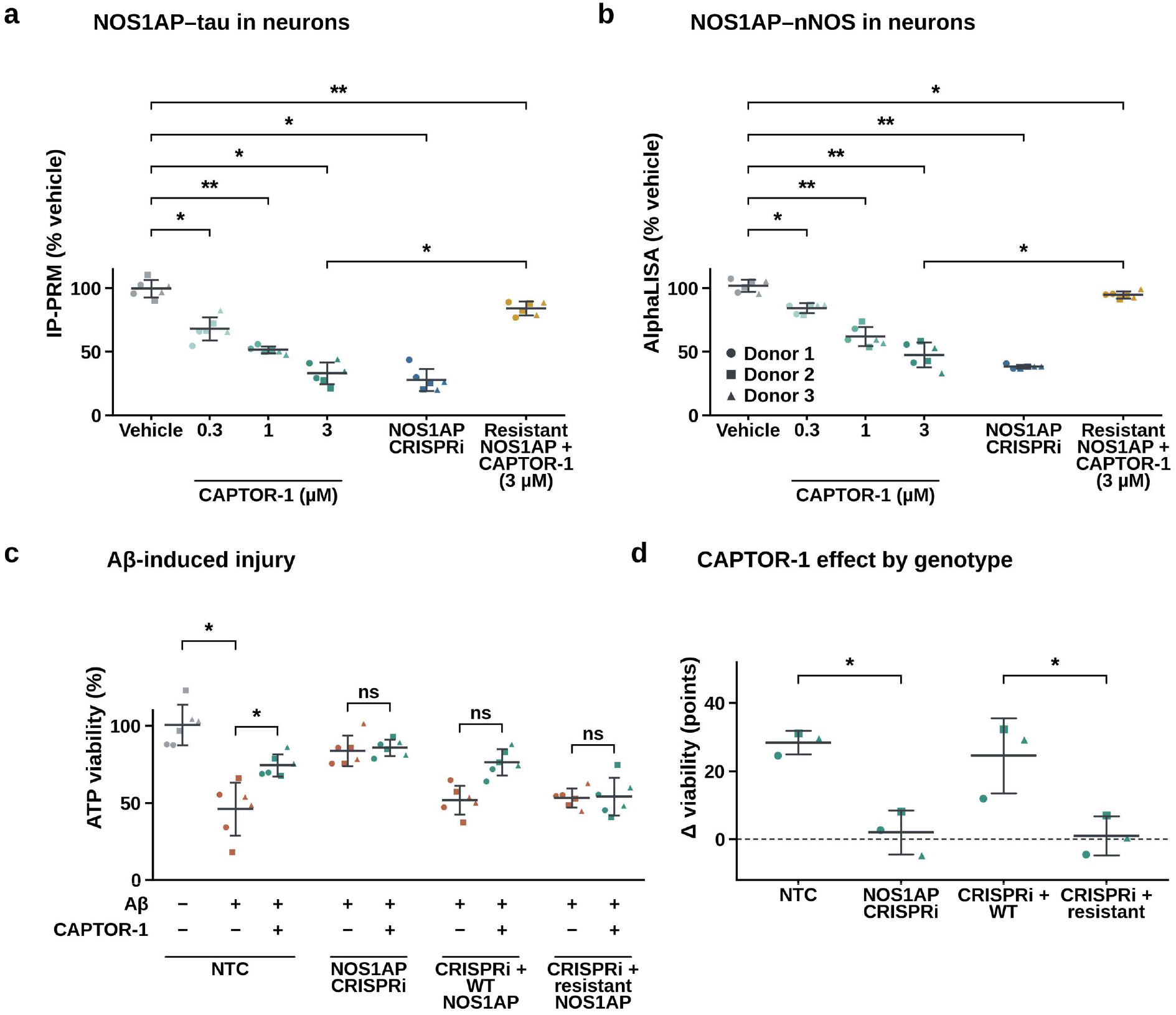
CAPTOR-1 acts through NOS1AP in human neurons. **a)** Endogenous NOS1AP-tau association (IP-PRM, % vehicle) in iPSC-derived neurons treated with vehicle or **CAPTOR-1**, after NOS1AP CRISPRi, or in CRISPRi neurons re-expressing the drug-resistant NOS1AP variant (Y16A/F42A) and treated with **CAPTOR-1** (3 µM). Points are observations (3 donors × 2 differentiations; symbols mark donors); bars are mean ± S.D.; brackets, paired t-tests on donor means (n = 3), Holm-adjusted across six comparisons (* *P* < 0.05, ** *P* < 0.01). **b)** Endogenous NOS1AP-nNOS association (AlphaLISA, % vehicle) under the same conditions. **c)** ATP viability of neurons exposed to Aβ, with non-targeting control (NTC) or NOS1AP CRISPRi, and CRISPRi neurons re-expressing wild-type or resistant NOS1AP, with or without **CAPTOR-1** (design matrix below the axis). Brackets as in a (Aβ effect, unadjusted; **CAPTOR-1** effects, Holm-adjusted across four genotypes; ns, not significant). **d) CAPTOR-1** effect on viability (Aβ + **CAPTOR-1** minus Aβ) per donor; brackets, paired t-test between genotypes, Holm-adjusted across two contrasts.

#### A drug-resistant NOS1AP variant restores the interactions

In NOS1AP CRISPRi neurons re-expressing the drug-resistant variant Y16A/F42A (Fig. 3b), **CAPTOR-1** (3 µM) left the NOS1AP-tau association at 84 ± 5% of vehicle-treated control neurons, compared with 33 ± 9% in control neurons treated with **CAPTOR-1** (Holm-adjusted *P* = 0.019; Fig. 5a), and the NOS1AP-nNOS association at 95 ± 3% (versus 48 ± 10%; *P* = 0.013; Fig. 5b). The residual reduction relative to vehicle (*P* = 0.002 and 0.048) is consistent with the weak but measurable affinity of the variant for **CAPTOR-1** (Kd 28 µM; Fig. 3b). The cellular effect of **CAPTOR-1** on both complexes therefore requires a drug-sensitive NOS1AP.

#### The neuroprotective effect of CAPTOR-1 depends on NOS1AP

We next asked whether the protective effect of **CAPTOR-1** in a disease-relevant insult depends on NOS1AP. Exposure of non-targeting control neurons to amyloid-β (Aβ) reduced ATP viability to 46 ± 17% of vehicle (*P* = 0.017) and increased nitrite and lactate dehydrogenase (LDH) release (Fig. 5c; Supplementary Fig. 8). **CAPTOR-1** improved viability by 28.4 ± 3.4 points (Holm-adjusted *P* = 0.019) and lowered nitrite by 9.9 ± 2.8 µM and LDH release by 4.3 ± 0.7-fold (Holm-adjusted *P* = 0.10 and 0.026). NOS1AP knockdown was itself protective (viability 84 ± 10%; *P* = 0.0006 versus control neurons under Aβ), and **CAPTOR-1** added nothing on top of knockdown (+2.0 ± 6.5 points). Re-expression of wild-type NOS1AP restored Aβ toxicity (52 ± 9%; *P* = 0.015 versus knockdown) and the response to **CAPTOR-1** (+24.5 ± 11.0 points, positive in all three donors; Holm-adjusted *P* = 0.18), whereas the drug-resistant Y16A/F42A variant restored toxicity (53 ± 6%; *P* = 0.010) but not the response to **CAPTOR-1** (+1.0 ± 5.8 points). The **CAPTOR-1** effect on viability therefore differed between genotypes (control versus knockdown, Holm-adjusted *P* = 0.044; wild-type versus resistant re-expression, *P* = 0.044; Fig. 5d). The corresponding genotype-dependent differences in nitrite and LDH release did not remain significant after correction for multiple comparisons (Supplementary Fig. 8). These data show that **CAPTOR-1** acts through NOS1AP in human neurons. Because **CAPTOR-1** reduced both the tau and the nNOS associations, however, they do not by themselves attribute the neuroprotection to the tau interface.

### CAPTOR-1 reduces tau seeding and pathogenic tau phosphorylation

#### CAPTOR-1 reduces tau seeding in human neurons through NOS1AP

We first asked whether **CAPTOR-1** reduces tau seeding, using recombinant P301S 2N4R tau PFFs (100 nM monomer equivalent for 72 h) in human NGN2-induced neurons; **CAPTOR-1** was added 2 h before PFF exposure and remained present throughout the experiment. PFF increased the tau aggregate signal (HTRF) from 1.05 ± 0.31 to 4.96 ± 0.40-fold (*P* = 0.006; n = 3 culture batches) and the tau seeding activity measured in a biosensor assay from 1.00 ± 0.21 to 5.06 ± 0.68-fold (*P* = 0.011; Fig. 6a,b). **CAPTOR-1** reduced both readouts dose-dependently (HTRF, 3.98 ± 0.26, 2.56 ± 0.17 and 1.81 ± 0.05-fold at 0.3, 1 and 3 µM; Holm-adjusted *P* = 0.045, 0.013 and 0.018; biosensor, 3.46 ± 0.54, 2.64 ± 0.17 and 1.94 ± 0.31-fold; *P* = 0.032, 0.038 and 0.038), to levels similar to those after NOS1AP CRISPRi (1.93 ± 0.32 and 1.97 ± 0.35-fold; P = 0.021 and 0.038). **CAPTOR-1** added no further reduction in NOS1AP CRISPRi neurons (HTRF, 1.80 ± 0.24 versus 1.93 ± 0.32-fold, *P* = 0.69; biosensor, 1.89 ± 0.09 versus 1.97 ± 0.35-fold, *P* = 0.17), consistent with action through NOS1AP. Total tau was unchanged by PFF, **CAPTOR-1** or knockdown (94-102% of control; Supplementary Fig. 9), so the reduction in seeding is not explained by lower tau levels.

#### CAPTOR-1 reduces pathogenic tau phosphorylation through NOS1AP

To test whether **CAPTOR-1** also lowers pathological tau phosphorylation, we performed quantitative phosphoproteomic profiling of NGN2-induced neurons from four neurologically normal donor lines, comparing unseeded cultures with matched cultures exposed to P301S tau PFFs, as described in Methods. PFF-exposed neurons showed increased phosphorylation of tau at T217, T231 and S202 (log2 fold versus control +1.37, +1.12 and +1.03; Holm-adjusted *P* = 0.002, 0.0005 and 0.0004; Fig. 6c-e; Supplementary Table 5). **CAPTOR-1** reduced these increases by 61%, 64% and 56% (to +0.54, +0.40 and +0.45; *P* = 0.006, 0.002 and 0.006), and NOS1AP CRISPRi reduced them similarly (+0.49, +0.38 and +0.28; *P* = 0.007, 0.007 and 0.003). In CRISPRi neurons re-expressing the drug-resistant NOS1AP variant Y16A/F42A, **CAPTOR-1** no longer lowered tau phosphorylation (+1.22, +1.02 and +0.84; *P* = 0.0003, 0.0007 and 0.008 versus **CAPTOR-1**-treated PFF-exposed neurons), which remained at 89%, 91% and 82% of the PFF-induced level.

#### CAPTOR-1 shifts associated signaling nodes toward control

In an exploratory analysis of nine further phosphosites, PFF-exposed neurons showed increased phosphorylation of GSK3β (Y216), p38α (T180/Y182), FYN (Y420), NOS1 (S1417) and GluN2B (Y1472) and decreased phosphorylation of CaMKIIα (T286), CREB (S133), SYN1 (S9) and PSD-95 (S73) (Fig. 6f; Supplementary Fig. 10). **CAPTOR-1** shifted all nine sites toward control, but with four donors and correction across nine sites only CREB pS133 differed significantly from untreated PFF-exposed neurons (Holm-adjusted *P* = 0.025). The resistant variant restored the PFF-exposed pattern (*P* < 0.05 versus **CAPTOR-1**-treated PFF-exposed neurons for GSK3β, p38α and CREB). Because **CAPTOR-1** also engages the NOS1AP-nNOS complex (Fig. 5b), these data show that the effects on tau seeding and phosphorylation depend on NOS1AP but do not assign them to the tau interface alone.

**Figure 6.**
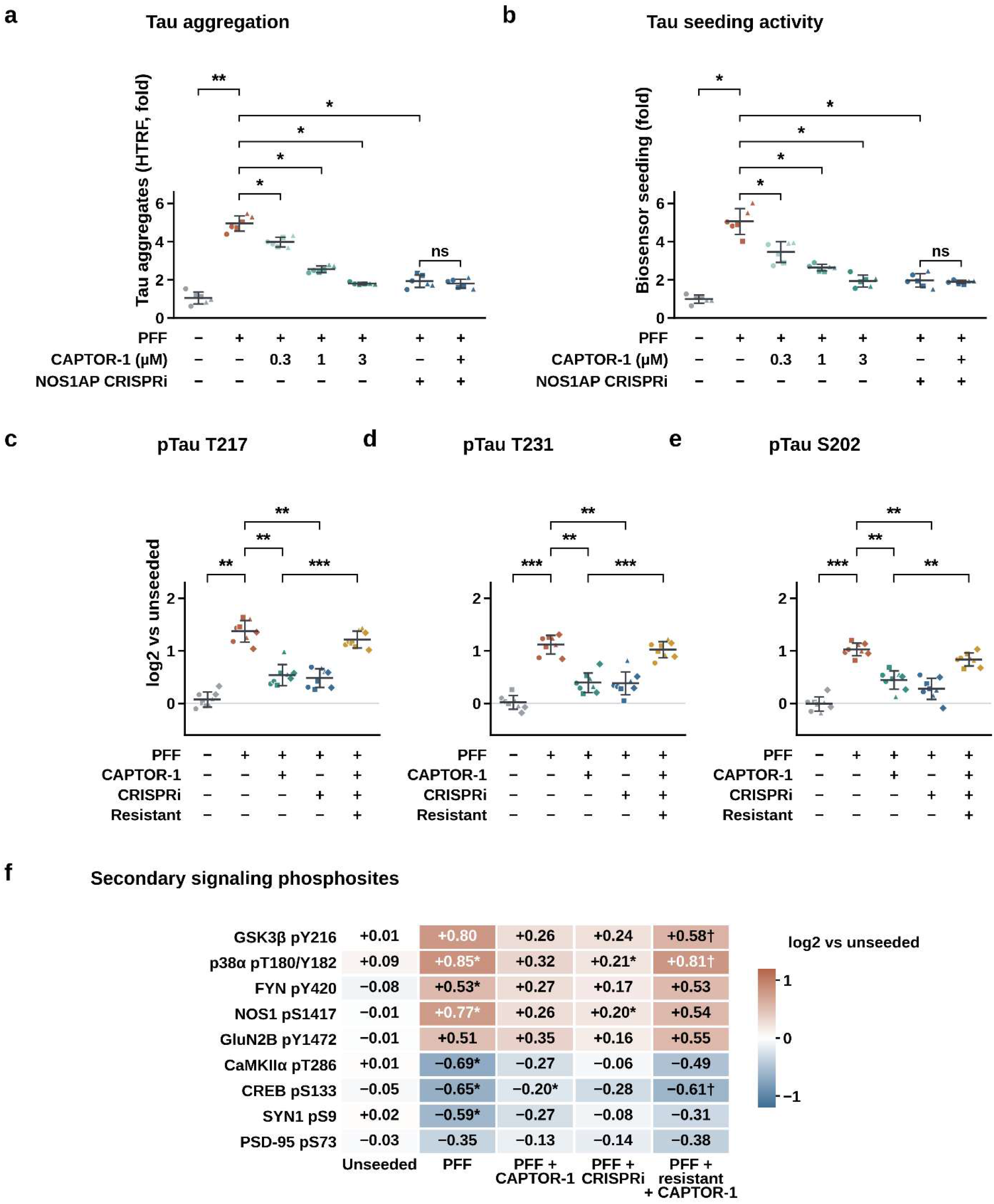
CAPTOR-1 reduces tau seeding and pathogenic tau phosphorylation. **a)** Tau aggregate signal (HTRF, fold) in human neurons seeded with PFF, treated with **CAPTOR-1**, after NOS1AP CRISPRi, or both (design matrix below the axis; the **CAPTOR-1** dose in the CRISPRi group is 3 µM). Points are observations (3 culture batches × 2 replicates; symbols mark batches); bars are mean ± S.D.; brackets, paired t-tests on batch means (n = 3). **CAPTOR-1** and CRISPRi versus PFF alone are Holm-adjusted across four comparisons; the PFF effect and the CRISPRi versus CRISPRi + **CAPTOR-1** contrast are single planned comparisons (* *P* < 0.05, ** *P* < 0.01; ns, not significant). **b)** Tau seeding activity (biosensor, fold) under the same conditions. **c-e)** Phosphorylation of tau at T217 **(c)**, T231 **(d)**, and S202 **(e)** in PFF-induced tau-pathology neurons, expressed as log2 fold change relative to matched unseeded controls: unseeded control, PFF, PFF + **CAPTOR-1**, PFF + NOS1AP CRISPRi, and PFF + CRISPRi + resistant NOS1AP (Y16A/F42A) + **CAPTOR-1**. Points are observations from four donor lines with two independent differentiations per line; symbols identify donors. Brackets indicate paired two-sided t-tests on donor-level means (n = 4), with Holm adjustment across the three prespecified tau sites. **f)** Mean phosphorylation of nine exploratory signaling sites, expressed as log2 fold change relative to matched unseeded controls. * *P* < 0.05 versus PFF-exposed neurons (PFF-exposed neurons column: versus control) and † *P* < 0.05 for resistant + **CAPTOR-1** versus PFF-exposed neurons + **CAPTOR-1**, Holm-adjusted across the nine sites.

#### CAPTOR-1 rescues NOS1AP-related deficits in patient-derived AD neurons

To extend our findings beyond the engineered CRISPRi and rescue models, we evaluated NGN2-induced neurons generated from eight commercially obtained Coriell iPSC donor lines: four neurologically normal controls and four patients with sporadic AD (see Methods). Each line underwent two independent differentiations, with the donor line used as the biological replicate. At baseline, AD-line neurons differed from control-line neurons on every readout: lower ATP viability (68 ± 2% versus 102 ± 3%, *P* < 0.0001), higher tau phosphorylation (pTau217, 95 ± 3 versus 26 ± 5 pg/mg, *P* < 0.0001), higher tau seeding activity (3.5 ± 0.1 versus 0.9 ± 0.1-fold, *P* < 0.0001), lower network burst activity (61 ± 7% versus 100 ± 7%, *P* = 0.0003) and higher nitrite (10.6 ± 0.5 versus 5.1 ± 0.5 µM, *P* < 0.0001) (Fig. 7a-e). **CAPTOR-1** improved all five readouts in AD-line neurons in a dose-dependent manner (Holm-adjusted P ≤ 0.02 for every readout at both 1 and 3 µM), whereas it had no significant effect on any readout in control-line neurons. These data show that the deficits captured by NOS1AP-dependent readouts in engineered cellular models (Figs. 5, 6) are also present in neurons derived from AD patients, and that **CAPTOR-1** corrects them.

**Figure 7.**
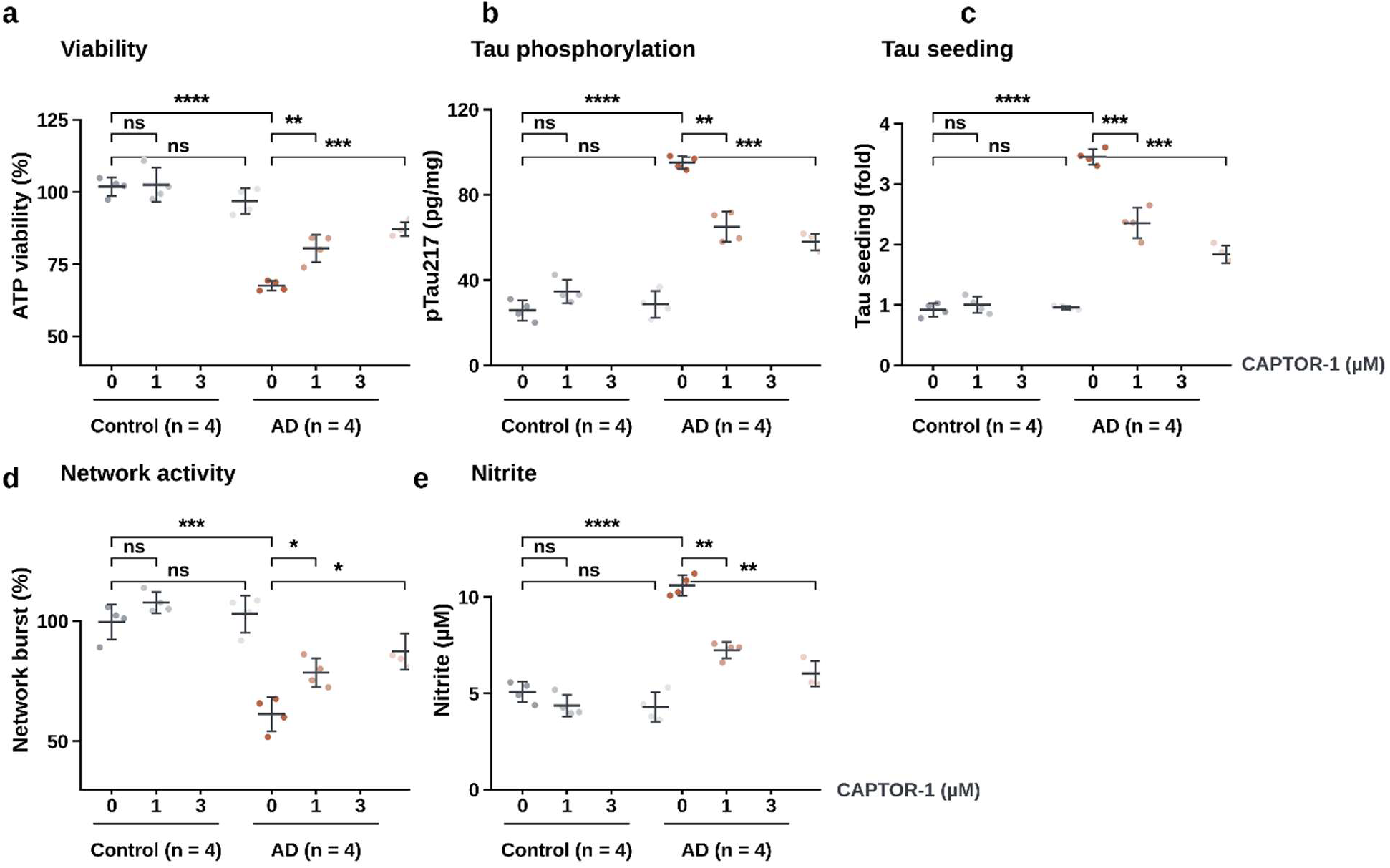
CAPTOR-1 rescues NOS1AP-related deficits in patient-derived AD neurons. **a-e)** ATP viability **(a)**, pTau217 **(b)**, tau seeding activity **(c)**, network burst activity **(d)** and nitrite **(e)** in iPSC-derived neurons from 4 control and 4 AD donor lines, treated with vehicle or **CAPTOR-1** (1 or 3 µM). Points are donor-level means (mean of 2 differentiations; n = 4 donors per diagnosis); bars are mean ± S.D. Brackets: baseline (vehicle) difference between diagnoses (unpaired t-test) and **CAPTOR-1** dose effect within each diagnosis (paired t-test on donor means, Holm-adjusted across 1 and 3 µM; **** *P* < 0.0001, *** *P* < 0.001, ** *P* < 0.01, * *P* < 0.05, ns not significant).

### Orally dosed CAPTOR-1 reaches the brain and engages NOS1AP in vivo

#### CAPTOR-1 has drug-like in vitro properties and is brain-penetrant

**CAPTOR-1** had adequate kinetic solubility (86 ± 9 µM at pH 7.4), moderate microsomal stability (mouse and human liver microsome t_1/2_, 54 ± 6 and 68 ± 8 min), high predicted passive permeability (PAMPA-BBB *P_e_*, 8.1 ± 0.9 × 10^−6^ cm/s) and low hERG inhibition (18 ± 4% at 10 µM) (Supplementary Table 6). After a single oral dose (30 mg/kg, C57BL/6J mice), **CAPTOR-1** reached a plasma C_max_ of 5,090 ± 721 ng/mL at a median T_max_ of 0.5 h and a brain C_max_ of 1,408 ± 231 ng/mL at 1.5 h (n = 4; Fig. 8a). Brain exposure relative to plasma increased over time (Kp,AUC_0-24_ = 0.46; brain:plasma ratio 0.15 at 15 min to 1.28 at 24 h; Fig. 8b), indicating that **CAPTOR-1** is retained in brain after plasma levels decline. Comparison of dose-normalized oral and intravenous plasma exposure yielded an absolute oral bioavailability of 40%.

**Figure 8.**
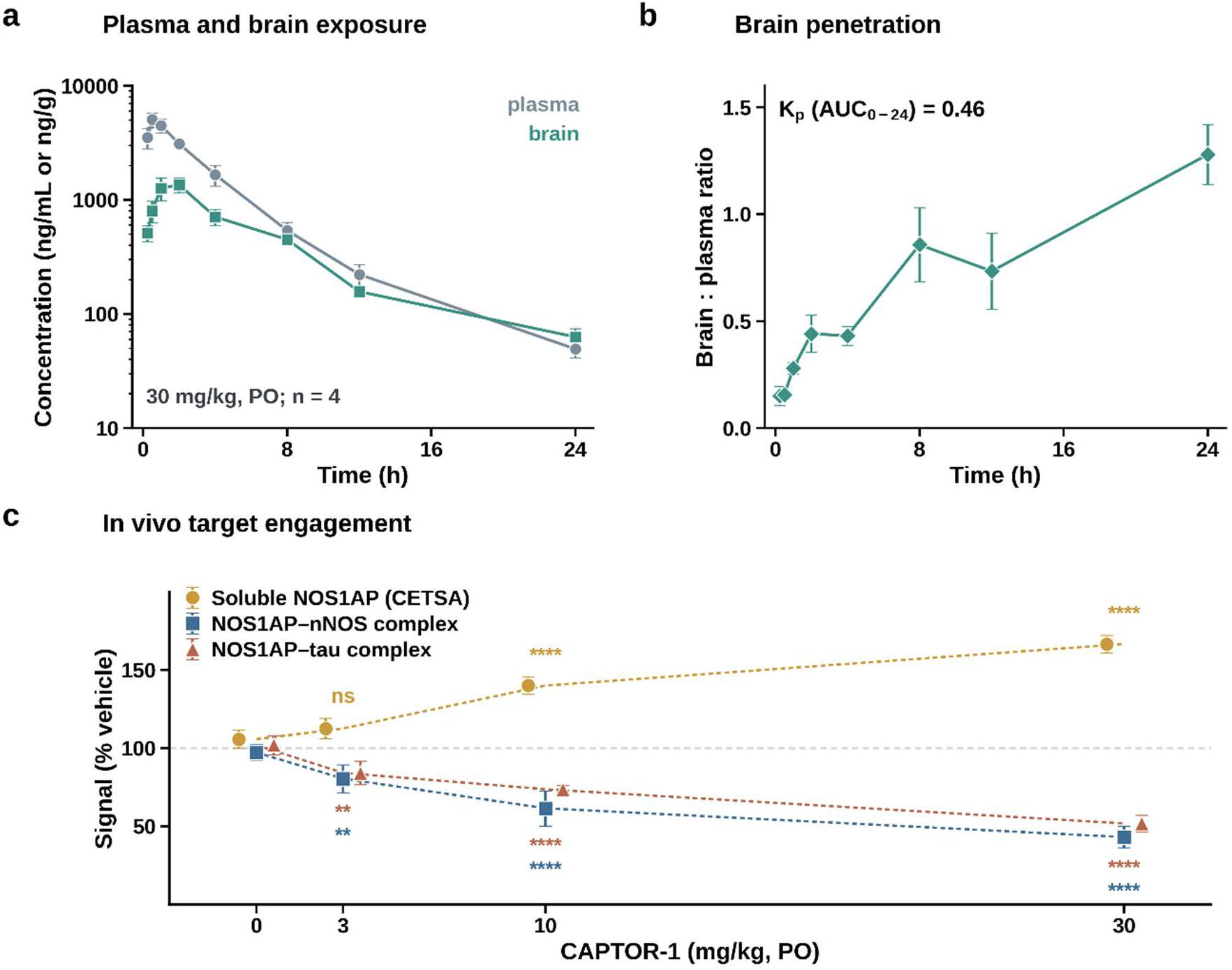
Orally dosed CAPTOR-1 reaches the brain and engages NOS1AP in vivo. **a)** Plasma and brain concentration of **CAPTOR-1** after a single oral dose (30 mg/kg); mean ± S.D., n = 4. **b)** Brain-to-plasma concentration ratios calculated from matched samples at each time point; Kp,AUC is the ratio of brain to plasma AUC_0-24_. **c)** In vivo target engagement: soluble NOS1AP after a thermal challenge (CETSA) and the NOS1AP-nNOS and NOS1AP-tau complexes (AlphaLISA) in brain homogenate, 2 h after a single oral dose of **CAPTOR-1** (mean ± S.D., n = 6 per dose). Stars: two-sided t-tests versus vehicle, Holm-adjusted across the three doses for each readout (** *P* < 0.01, **** *P* < 0.0001, ns not significant).

#### Orally dosed CAPTOR-1 engages NOS1AP in vivo

Before initiating the in vivo studies, we confirmed that **CAPTOR-1** retained cross-species target binding, with affinities of 0.22 ± 0.04 µM by MST and 0.28 ± 0.05 µM by BLI for recombinant mouse NOS1AP (mean ± S.D., n = 3 independent experiments).

To confirm target engagement in vivo, adult C57BL/6J mice, 10-12 weeks old and balanced by sex, received vehicle or a single oral dose of **CAPTOR-1** at 3, 10, or 30 mg/kg; brain tissue was collected 2 h after dosing, and NOS1AP thermal stabilization and the NOS1AP-nNOS and NOS1AP-tau complexes were measured in brain homogenates (n = 6 per dose). Soluble NOS1AP after the heat challenge increased dose-dependently (113 ± 7%, 140 ± 6% and 167 ± 6% of vehicle at 3, 10 and 30 mg/kg; Holm-adjusted *P* = 0.084, < 0.0001 and < 0.0001; Fig. 8c), consistent with target engagement. In the same brain homogenates, the NOS1AP-nNOS complex fell to 81 ± 9%, 62 ± 11% and 43 ± 7% of vehicle (*P* = 0.002, < 0.0001 and < 0.0001) and the NOS1AP-tau complex to 84 ± 8%, 74 ± 3% and 52 ± 5% (*P* = 0.001, < 0.0001 and < 0.0001).

Both complexes were reduced at every dose tested, so, as in cultured neurons (Fig. 5b) and patient-derived neurons (Fig. 7), **CAPTOR-1** engages NOS1AP at both interfaces in vivo, with a numerically larger effect on the tau complex at the two higher doses.

### CAPTOR-1 rescues tau pathology, synaptic function and memory in a tauopathy model (PS19)

#### Chronic CAPTOR-1 reduces tau pathology and rescues synaptic and cognitive deficits in PS19 mice

To test efficacy in vivo, we dosed PS19 mice and wild-type (WT) littermates orally with **CAPTOR-1** or vehicle (30 mg/kg once daily, the dose used for target engagement) from 6 to 8 months of age (n = 16 per group, 8 females and 8 males; Fig. 9). At 8 months, vehicle-treated PS19 mice showed the expected tauopathy phenotype relative to WT: reduced contextual fear memory (24 ± 5% versus 51 ± 6% freezing), impaired hippocampal long-term potentiation (103 ± 10% versus 157 ± 15% fEPSP), and elevated tau aggregation, phosphorylation (pTau181, pTau231), seeding activity and CSF neurofilament light (NfL) (all Holm-adjusted *P* < 0.0001). Chronic **CAPTOR-1** improved every readout in PS19 mice (freezing 38 ± 6%, LTP 137 ± 17%, and 45-63% reductions in the five molecular tau and neurodegeneration readouts; all Holm-adjusted *P* < 0.0001 versus PS19-vehicle), without recovering fully to WT levels. **CAPTOR-1** had no effect on any readout in WT mice (Holm-adjusted *P* = 1.00 for freezing, LTP, pTau181 and tau seeding), and a genotype × treatment interaction confirmed that the drug effect was specific to PS19 mice for every readout (*P* < 0.0001, except freezing *P* = 0.0001).

**Figure 9.**
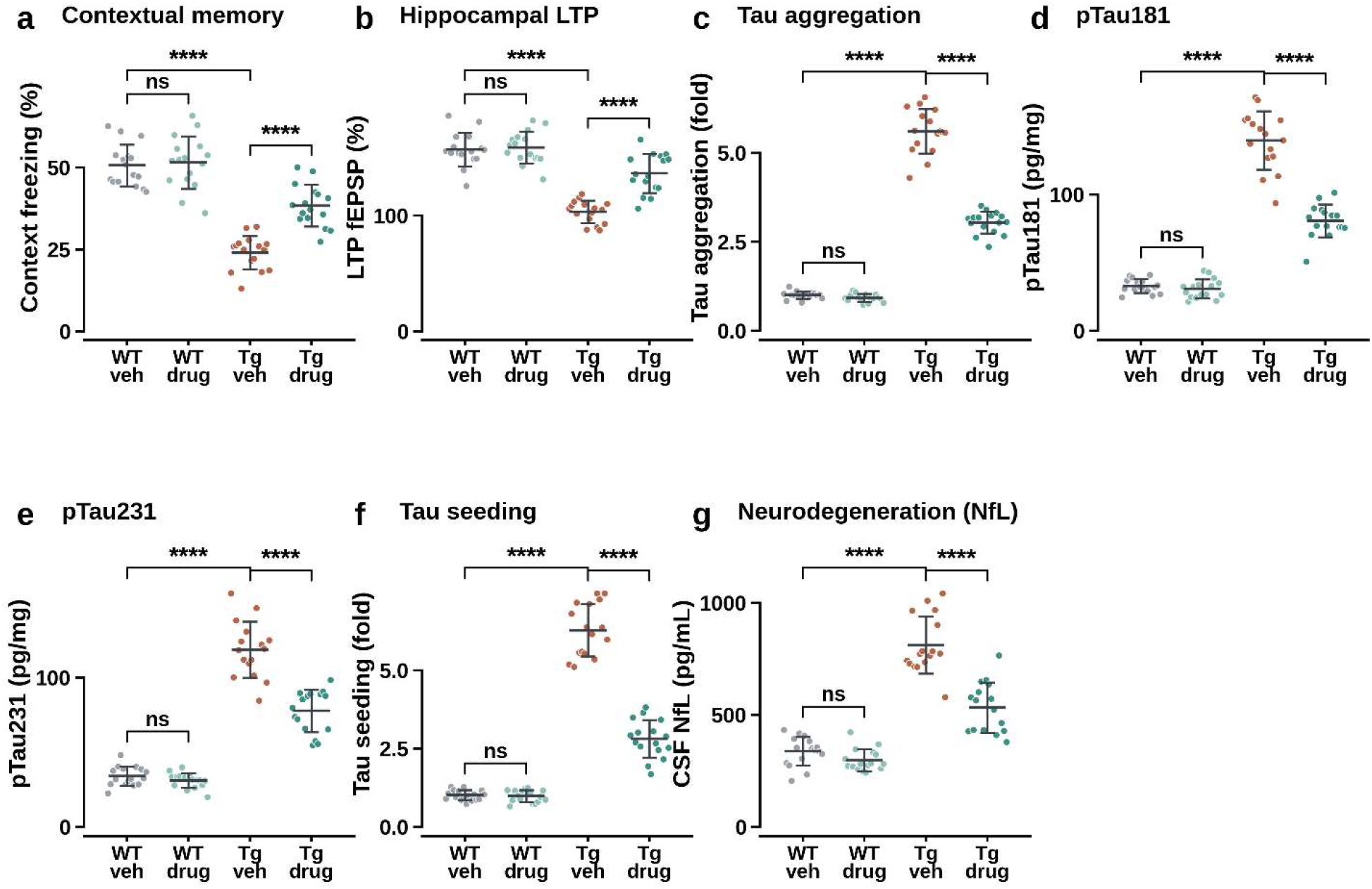
CAPTOR-1 rescues tau pathology, synaptic function and memory in PS19 mice. **a)** Contextual fear memory (freezing, 24 h post-conditioning). **b)** Hippocampal CA1 long-term potentiation (fEPSP, 50-60 min post-theta-burst stimulation). **c-f)** Hippocampal tau aggregation (HTRF) **(c)**, pTau181 **(d)**, pTau231 **(e)** and tau seeding activity **(f)**. **g)** CSF neurofilament light (NfL). Points are individual animals (n = 16 per group, 8 females and 8 males); bars are mean ± S.D. Brackets: disease effect (WT-vehicle versus PS19-vehicle) and **CAPTOR-1** effect within each genotype (Welch t-tests, Holm-adjusted across the 7 readouts within each contrast family; **** *P* < 0.0001, ns not significant). Both PS19 groups remained significantly different from WT-vehicle for every readout (adjusted *P* < 0.05; comparisons not shown).

### CAPTOR-1 rescues amyloid-associated deficits in an amyloidosis model with secondary tau pathology (APP/PS1)

#### Chronic CAPTOR-1 reduces amyloid pathology and rescues memory and synaptic deficits in APP/PS1 mice

To test whether efficacy extends to an amyloidosis model with secondary tau pathology, we dosed APP/PS1 mice and WT littermates as for PS19 (30 mg/kg once daily, oral, 8 to 10 months of age; n = 16 per group; Fig. 10). Vehicle-treated APP/PS1 mice showed impaired spatial memory (RAWM errors, 7.9 ± 1.3 versus 3.2 ± 0.7 in WT), reduced contextual freezing (21 ± 8% versus 46 ± 7%), impaired LTP (111 ± 12% versus 161 ± 12% fEPSP), and elevated brain Aβ_42_, CSF NfL and cortical nitrite (all Holm-adjusted *P* < 0.0001). Chronic **CAPTOR-1** improved every readout in APP/PS1 mice (RAWM errors 4.7 ± 0.9, freezing 41 ± 6%, LTP 143 ± 18%, and 31-45% reductions in Aβ_42_, NfL and nitrite; all Holm-adjusted *P* < 0.0001 versus APP/PS1-vehicle). **CAPTOR-1** had no significant effect in WT mice after correction (Holm-adjusted P = 1.00 for RAWM, LTP, Aβ_42_ and NfL; *P* = 0.09 for freezing), and a genotype × treatment interaction confirmed a genotype-specific drug effect for every readout (*P* ≤ 0.0006). The data from both animal models show that chronic oral **CAPTOR-1** rescues pathology, synaptic function and cognition in both a primary tauopathy model and an amyloidosis model with secondary tau pathology.

**Figure 10.**
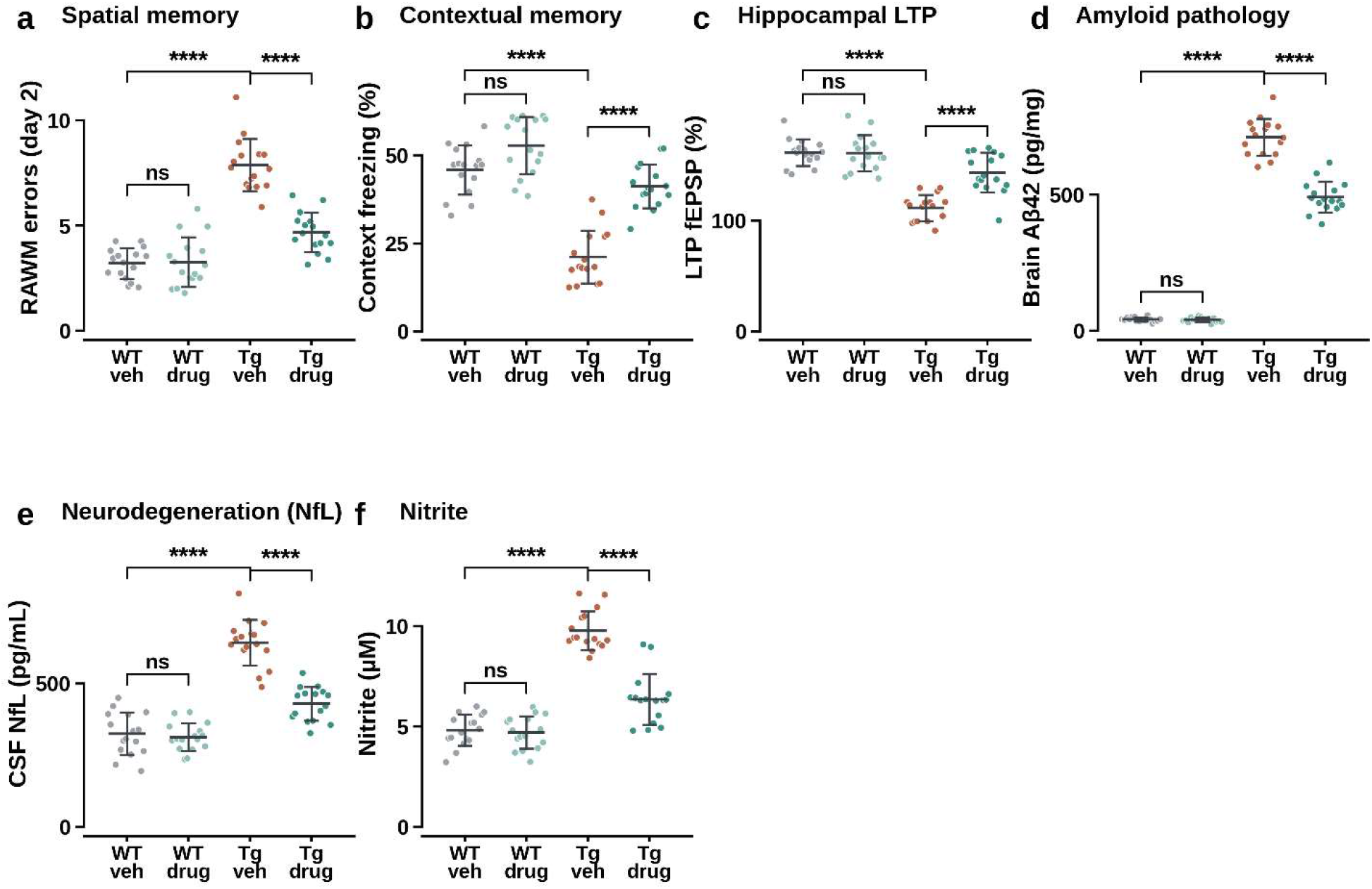
CAPTOR-1 rescues amyloid-associated deficits in APP/PS1 mice. **a)** Spatial memory (radial-arm water maze, errors on day 2). **b)** Contextual fear memory (freezing, 24 h post-conditioning). **c)** Hippocampal CA1 long-term potentiation (fEPSP). **d)** Brain Aβ_42_ (guanidine-extractable, cortical homogenate). **e)** CSF neurofilament light (NfL). **f)** Cortical nitrite. Points are individual animals (n = 16 per group, 8 females and 8 males); bars are mean ± S.D. Brackets: disease effect (WT-vehicle versus APP/PS1-vehicle) and **CAPTOR-1** effect within each genotype (Welch t-tests, Holm-adjusted across the 6 readouts within each contrast family; **** *P* < 0.0001, ns not significant).

## DISCUSSION

Here we report **CAPTOR-1**, to our knowledge the first small molecule that directly disrupts NOS1AP-tau interaction. Unlike the compounds previously developed against the NOS1AP-nNOS interface, including the anxiolytic ZLc-002 [15], **CAPTOR-1** was discovered by a screen against full-length NOS1AP and engages a pocket in the protein’s N-terminal region that overlaps the tau-binding surface (Figs. 2-4). It thereby converts a genetically validated but previously untargeted node in tau biology into a chemical starting point for drug discovery.

Three independent lines of evidence tie cellular and in vivo effects of **CAPTOR-1** to NOS1AP itself rather than to an off-target action. Substitution of the two pocket residues substantially attenuated **CAPTOR-1** activity across the principal mechanistic and cellular readouts, including NOS1AP-tau complex disruption, ATP viability, tau phosphorylation, and tau-seeding activity, whereas the corresponding genotype-dependent differences in nitrite and LDH release did not remain significant after correction for multiple comparisons. Genetic knockdown of NOS1AP by CRISPR interference phenocopied **CAPTOR-1** at every readout, with no additive effect when the two were combined (Figs. 5, 6). And the compound was inactive against 20 structurally related and unrelated proteins, including tau itself (Fig. 2e). At the same time, **CAPTOR-1** reduces the NOS1AP-nNOS complex in cells and in vivo, albeit less than the NOS1AP-tau complex at matched concentrations (Figs. 4d, 5b, 8c). The pocket therefore sits close enough to both interfaces that engaging it perturbs each, and the selectivity we observe is a difference of degree rather than of kind. Structural resolution of the pocket, which currently rests on mutagenesis, HDX-MS and homology modeling rather than a solved structure (Fig. 3), together with a variant that separates the two interfaces more completely than Y16A/F42A, would be needed to establish how much of the remaining nNOS engagement is intrinsic to this pocket and how much could still be engineered away.

A second aim of this study was to test how far the phenotype generalizes beyond a single engineered system. The direction, and in most cases the approximate magnitude, of the **CAPTOR-1** effect was preserved across isogenic knockdown-and-rescue neurons (Figs. 5, 6), commercially sourced neurons from AD patients (Fig. 7a-e), and two independent mouse models representing a primary tauopathy (PS19) and an amyloidosis with secondary tau pathology (APP/PS1) (Figs. 9, 10). Several of the readouts used to track this effect, tau phosphorylated at T217 and CSF neurofilament light in particular, are also used as fluid biomarkers in AD clinical trials, which should make a biomarker-anchored proof-of-mechanism study in patients more tractable than for a target without such precedent.

The pharmacological properties of **CAPTOR-1** support this path. It is orally bioavailable, crosses the blood-brain barrier, and is retained in brain after plasma concentrations decline (Fig. 8a,b), and a single oral dose engages NOS1AP in brain in a dose-dependent manner (Fig. 8c). Eight weeks of once-daily oral dosing at this same dose reduced tau pathology, restored synaptic function, and improved memory in both mouse models without a detectable effect in wild-type littermates (Figs. 9, 10), arguing that the benefit requires the disease genotype rather than reflecting a nonspecific behavioral effect of the compound.

These results add a small molecule option to a field of tau-lowering approaches so far dominated by oligonucleotides and antibodies. MAPT-targeting antisense oligonucleotides markedly lower CSF tau [4], and anti-tau antibodies such as semorinemab engage tau in patients [5], but both require intrathecal or intravenous administration and act on tau itself rather than on an upstream regulator of its pathological modification. **CAPTOR-1** instead targets an adaptor protein genetically upstream of tau hyperphosphorylation and aggregation [16], is orally dosed, and, because it also modulates the NOS1AP-nNOS axis, may in principle combine some of the reported benefits of nNOS-interface-directed compounds [15] with a tau-directed disease-modifying effect; whether these two actions are complementary or simply redundant in vivo is not addressed by the present data.

Future work should resolve the structural basis of the NOS1AP pocket, test whether selectivity for the tau interface can be further improved, establish a dose-response relationship for chronic efficacy, and determine whether the benefit of **CAPTOR-1** is sustained after treatment withdrawal or requires combination with amyloid-or tau-directed therapies. With these caveats, the data presented here establish pharmacological disruption of the NOS1AP-tau interaction as a viable, disease-modifying strategy across purified protein, human cells, and two mouse models of AD, and provide **CAPTOR-1** as the chemical tool that makes further mechanistic and translational work on this target possible.

## MATERIALS AND METHODS

### Compounds and preparation

**NTB24**, **CAPTOR-1**, and the focused analog panel were obtained as analytically characterized dry powders from Molport. Compound identity was confirmed by liquid chromatography-mass spectrometry and purity was at least 95% by analytical high-performance liquid chromatography. Stock solutions were prepared at 10 or 20 mM in anhydrous dimethyl sulfoxide (DMSO), divided into single-use aliquots, and stored at −80°C. Working solutions were prepared immediately before use. DMSO was matched across conditions and did not exceed 2.5% in biophysical screening, 1% in purified-protein assays, or 0.1% in cell-based assays. **CAPTOR-1** was formulated for oral dosing in 0.5% methylcellulose and 0.1% Tween-80 in purified water.

### Recombinant proteins

Human full-length NOS1AP (UniProt O75052; residues 1-503) and NOS1AP fragments were expressed in Escherichia coli BL21(DE3) with an N-terminal hexahistidine tag followed by a tobacco etch virus protease cleavage site. Cultures were induced with 0.4 mM isopropyl beta-D-1-thiogalactopyranoside at 18°C for 16 h. Cell pellets were disrupted by sonication in 50 mM Tris-HCl pH 7.5, 300 mM NaCl, 10 mM imidazole, 1 mM tris(2-carboxyethyl)phosphine, and protease inhibitors. Clarified lysates were purified by nickel-affinity chromatography, and proteins were polished by size-exclusion chromatography on a Superdex 200 Increase column in 20 mM HEPES pH 7.4, 150 mM NaCl, and 1 mM tris(2-carboxyethyl)phosphine. Full-length human 2N4R tau and tau fragments were produced by bacterial expression, heat clarification, cation-exchange chromatography, and size-exclusion chromatography. Purity exceeded 90% by capillary electrophoresis, and monodispersity was verified by analytical size-exclusion chromatography. Protein concentrations were determined from absorbance at 280 nm using sequence-derived extinction coefficients.

### Affinity selection mass spectrometry screening

A 10,000-compound drug-like HitFinder collection (ThermoFisher Scientific, Catalog #HF13384) was screened against full-length NOS1AP by affinity selection-mass spectrometry (AS-MS), following our previously reported solution-phase workflows [26]. Compounds were arrayed in pools of 100 at 1 µM per compound and incubated with 2 µM NOS1AP for 60 min at 25°C in 20 mM HEPES pH 7.4, 150 mM NaCl, 0.01% Tween-20, and 1% DMSO. Matched protein-free and heat-denatured-protein controls were processed in parallel. Protein-ligand complexes were separated from unbound compounds by rapid size-exclusion chromatography. Bound molecules were released with 80% methanol containing internal standards and identified by high-resolution liquid chromatography-mass spectrometry using retention time and accurate mass. Enrichment was calculated relative to both controls. Compounds enriched at least threefold with reproducible ion direction in duplicate pools were considered primary hits. Reactive structures, unstable compounds, duplicates, and compounds with ambiguous mass-spectral assignments were removed by manual triage. The 50 remaining candidates were advanced to single-concentration MST validation.

### Microscale thermophoresis

Binding of **NTB24** and analogs to NOS1AP was measured on a Monolith X instrument (NanoTemper Technologies). His-tagged full-length NOS1AP was labeled with RED-tris-NTA dye according to the manufacturer’s instructions and used at 20 nM in phosphate-buffered saline containing 0.05% Tween-20 and 2.5% DMSO. Compounds were prepared as 16-point twofold serial dilutions. Labeled protein and compound were incubated for 20 min at room temperature before loading into premium-coated capillaries. Measurements were acquired at 25°C with medium infrared-laser power and 20% excitation power. Fluorescence changes and capillary scans were inspected for adsorption, aggregation, or compound fluorescence. Three technical measurements were acquired at each concentration. Curves were fit by nonlinear least squares to a 1:1 binding isotherm using the total ligand concentration. The fitted dissociation constant and its standard error are reported. Curves that did not reach a plateau or produced poorly constrained confidence intervals were reported as Kd greater than 30 µM rather than assigned a point estimate. Binding of NOS1AP to tau was measured independently using three protein preparations, with fluorescently labeled NOS1AP and unlabeled 2N4R tau over 0.01-10 µM; the mean Kd and standard deviation across preparations are reported.

### Protein selectivity panel

**CAPTOR-1** binding was profiled by biolayer interferometry (BLI) on an Octet RED96e instrument against full-length NOS1AP and 20 comparator proteins: the nNOS PDZ domain, PSD-95, PSD-93, SAP97, SAP102, α1-syntrophin, SAPAP1, MAGI2, SHANK3, full-length 2N4R tau, MAP2, MAP1B, FYN SH2, APOE4, TREM2, 14-3-3ζ, GAPDH, carbonic anhydrase II, bovine serum albumin, and human serum albumin. Biotinylated proteins were immobilized on streptavidin biosensors to matched loading responses. **CAPTOR-1** was initially tested at 10 µM using five independently loaded sensors per protein. Tau, the nNOS PDZ domain, PSD-95, and PSD-93 were additionally evaluated across a 0.3-30 µM **CAPTOR-1** concentration series. Responses were reference-and background-subtracted and expressed relative to the maximal NOS1AP response obtained in the same experiment.

### Biolayer interferometry

Direct binding of **CAPTOR-1** to NOS1AP was measured by biolayer interferometry on an Octet RED96e instrument at 25°C with orbital shaking at 1,000 rpm. His-tagged full-length NOS1AP, 2N4R tau, or tag-only control protein was loaded onto separate Ni-NTA biosensors at 10 micrograms mL^−1^ for 120 s to a loading response of 0.8-1.2 nm. Sensors were equilibrated in phosphate-buffered saline pH 7.4 containing 0.01% Tween-20, 0.1% bovine serum albumin, and 1% DMSO. After a 60-s baseline, sensors were transferred to **CAPTOR-1** for 120 s and then to compound-free buffer for 300 s. A twofold dilution series from 0.03 to 30 µM was tested. Reference-sensor and buffer-only signals were subtracted, and solvent correction was applied. Equilibrium responses were fit to a 1:1 binding model. Four independent sensor loadings were performed with freshly prepared sensors and analyte series. Each loading yielded an independent Kd; the reported affinity is the mean and standard deviation of these four values.

### NOS1AP mutagenesis and protein-quality controls

NOS1AP Y16F, Y16A, F42A, and Y16A/F42A variants were generated in the full-length NOS1AP expression plasmid by site-directed mutagenesis and verified by complete insert sequencing. Each variant was expressed and purified independently. **CAPTOR-1** affinity was measured by BLI as described above. Analytical size-exclusion chromatography was performed on a Superdex 200 Increase 10/300 GL column; monomer content was calculated from the monomer peak area divided by the total integrated protein area. Apparent melting temperatures were determined by differential scanning fluorimetry using SYPRO Orange. Cellular expression was measured after transient expression in HEK293T cells by targeted parallel-reaction-monitoring mass spectrometry normalized to a coexpressed reference protein. Basal NOS1AP-tau association was measured by AlphaLISA in the absence of compound.

### Hydrogen deuterium exchange mass spectrometry

Hydrogen-deuterium exchange mass spectrometry (HDX-MS) was performed with purified full-length NOS1AP. Protein at 10 µM was incubated for 30 min at 25 degrees C with vehicle or 100 µM **CAPTOR-1**, giving a 1:10 protein-to-ligand molar ratio. Exchange was initiated by tenfold dilution into the corresponding deuterated buffer and allowed to proceed for 30, 300, or 3,000 s at 20°C. Reactions were quenched to pH 2.5 at 0°C, digested online with immobilized pepsin, trapped, and separated at 0°C by reversed-phase ultraperformance liquid chromatography before high-resolution mass spectrometry. Peptides were identified from undeuterated controls with a 1% false-discovery rate. The peptide map covered 82% of the NOS1AP sequence. Deuterium uptake was corrected using undeuterated and maximally deuterated controls. Three independently prepared labeling reactions were analyzed per condition and time point. A peptide was considered ligand responsive when the absolute uptake difference exceeded 0.5% and the two-sided peptide-level comparison was significant after false-discovery-rate correction across the peptide map. The eight peptides presented in the supplementary data are all peptides meeting these criteria, not a selected subset.

### Molecular docking and dynamics simulations

Ligand geometries were built in ChemBio3D Ultra 12.0, preoptimized with the MM2 force field, and further optimized in Gaussian 09 at the B3LYP/6-31G(d) level. Docking used the published NOS1AP N-terminal homology model [26], restricted to residues 1-160. The receptor was prepared by removing nonprotein atoms and water, adding polar hydrogen atoms, and assigning Gasteiger charges. AutoDock 4.2 was used with a 40 x 40 x 40 point grid, 0.375-Å spacing, centered on the predicted N-terminal pocket. The Lamarckian genetic algorithm was run independently 100 times for each ligand; the lowest-energy member of the largest cluster was advanced to molecular dynamics.

The apo NOS1AP model and complexes with **NTB24** or **CAPTOR-1** were simulated in AMBER 20. Protein and ligand parameters were assigned with ff14SB and GAFF2, respectively, and ligand charges were derived by the AM1-BCC procedure. Each system was solvated in a truncated-octahedral TIP3P water box extending at least 15 Å from the solute and neutralized with counterions. Systems underwent 5,000 steps each of restrained steepest-descent and conjugate-gradient minimization, followed by the same unrestrained sequence. After gradual heating from 0 to 300 K, a 100-ns production trajectory was collected in the NPT ensemble at 300 K and 1 bar with a 2-fs time step. Hydrogen-containing bonds were constrained with SHAKE, the nonbonded cutoff was 10 Å, and long-range electrostatics were treated by particle-mesh Ewald. Coordinates were saved every 10 ps. One independent trajectory was generated per system. Root-mean-square deviation, radius of gyration, and residue-level root-mean-square fluctuation were calculated after alignment to the protein backbone.

Protein-ligand contacts were calculated from the equilibrated 80-100 ns interval. A residue was defined as contacting the ligand when any protein and ligand heavy atoms were within 4.0 Å; persistent contacts were present in at least 30% of analyzed frames. Binding energies were estimated with the molecular mechanics/Poisson-Boltzmann surface area method in MMPBSA.py. Forty-one snapshots were selected every 50 frames from 80 to 100 ns. Reported values include van der Waals, electrostatic, polar-solvation, and nonpolar-solvation terms. Conformational entropy was not calculated. Because each system was represented by one trajectory and adjacent snapshots were correlated, the calculations were used only to compare the direction of change from **NTB24** to **CAPTOR-1** and not as experimental affinity estimates.

### Surface plasmon resonance and interface mapping

NOS1AP-tau binding and construct mapping were measured on a Biacore T200 at 25°C in HBS-EP+ buffer. Full-length 2N4R tau or tau fragments were immobilized on CM5 sensor chips by amine coupling at low density; a mock-coupled flow cell served as reference. Full-length NOS1AP was injected from 0.03 to 2 µM for 120 s at 30 microliters min^−1^ followed by 300 s dissociation. Surfaces were regenerated with a short pulse of 10 mM glycine pH 2.5. Double-referenced sensorgrams were globally fit to a 1:1 kinetic model to obtain association and dissociation rate constants. Three independently prepared surfaces were analyzed. For interface mapping, NOS1AP fragments 1-75, 1-126, 1-150, 76-150, 127-503, and 151-503 and tau fragments comprising the N-terminal, proline-rich (151-243), microtubule-binding, and C-terminal regions were tested at matched concentrations. Responses were normalized to the full-length NOS1AP-full-length tau pair.

### Purified NOS1AP tau disruption assay

Disruption of the purified NOS1AP-tau complex was quantified by AlphaLISA in white 384-well plates at a final assay volume of 20 µL per well. His-tagged full-length NOS1AP and biotinylated full-length 2N4R tau were each used at 10 nM in assay buffer containing 25 mM HEPES (pH 7.4), 150 mM NaCl, 0.1% bovine serum albumin, and 0.01% Tween-20. **CAPTOR-1** was tested as a 12-point threefold dilution with a final DMSO concentration of 0.5%. **CAPTOR-1** was preincubated with NOS1AP for 30 min at room temperature, followed by addition of tau and incubation for 60 min. AlphaLISA Nickel Chelate Acceptor Beads (AL108C; Revvity) were added to a final concentration of 20 µg/mL, followed 30 min later by Alpha Streptavidin Donor Beads (6760002; Revvity) at 20 µg/mL under subdued light. Plates were incubated for an additional 60 min at room temperature, and luminescence was measured using Tecan Spark plate reader. Four independent protein preparations were analyzed, with technical duplicates averaged within each experiment. Concentration-response data were fitted using four-parameter logistic regression. Assay-interference controls included 0.01% and 0.05% Tween-20, 0.5% and 1% DMSO, omission of either binding partner, a His-tag-only protein, and an unrelated intrinsically disordered protein. **CAPTOR-1** activity was interpreted only when the compound did not alter the corresponding background signal.

### Live cell NOS1AP nNOS counter screen

The NOS1AP-nNOS interaction was measured with the previously validated NanoBiT complementation assay [25]. CHO-K1 cells were transiently cotransfected with LgBiT-NOS1AP and HiBiT-nNOS constructs. Twenty-four hours later, cells were treated for 5 h with vehicle, NTB24, or **CAPTOR-1** over the stated concentration series. Nano-Glo Live Cell substrate was added and luminescence was recorded on a Tecan Spark plate reader. Signal was normalized to vehicle-treated cells on the same plate. Six wells were tested per concentration in one counter-screening experiment.

### Human iPSC derived cortical neurons

Human iPSC lines (from the Coriell Institute) were expanded under feeder-free conditions and differentiated into excitatory cortical neurons by doxycycline-inducible NGN2 expression. Cells were plated on tissue-culture plates precoated overnight at 4°C with poly-D-lysine hydrobromide (50 µg/mL; Sigma-Aldrich, P6407), followed by recombinant human laminin-521 (5 µg/mL; BioLamina, LN521) for 2 h at 37°C. Neurons were maintained in Neurobasal Plus Medium (Thermo Fisher Scientific, A3582901) supplemented with 2% B-27 Plus Supplement (Thermo Fisher Scientific, A3582801), 1% GlutaMAX Supplement (Thermo Fisher Scientific, 35050061), recombinant human brain-derived neurotrophic factor (BDNF; 10 ng/mL; PeproTech, 450-02), recombinant human neurotrophin-3 (NT-3; 10 ng/mL; PeproTech, 450-03), and doxycycline hyclate (2 µg/mL; Sigma-Aldrich, D9891). Half of the medium was replaced every 2-3 days. On day 3, neurons were replated with primary human astrocytes at a 4:1 neuron-to-astrocyte ratio. Experiments were performed on differentiation days 21-28. Each independent differentiation was initiated from a separate iPSC passage. Cultures were screened for mycoplasma before use. Neuronal identity and culture quality were confirmed by flow cytometry for MAP2 and betaIII-tubulin and by RT-qPCR for MAP2, RBFOX3, and SLC17A7; batches with viability below 85% or neuronal-marker positivity below 85% were excluded before treatment assignment.

### CRISPR interference and rescue

NOS1AP was depleted using iPSC lines stably expressing nuclease-inactive Cas9 fused to the KRAB transcriptional-repression domain (dCas9-KRAB). Two independent sgRNAs targeting sequences within 200 bp of the NOS1AP transcription start site were cloned individually into lentiviral sgRNA vectors and introduced by transduction at a multiplicity of infection below 1. A non-targeting sgRNA with no predicted target in the human genome served as the negative control. Transduced cells were selected with puromycin (1 µg/mL for 72 h) before NGN2 induction. NOS1AP knockdown was quantified independently by RT-qPCR, normalized to GAPDH and RPLP0 using the ΔΔCt method, and by targeted parallel-reaction-monitoring mass spectrometry using NOS1AP-specific proteotypic peptides. Differentiations were included only when both sgRNAs produced at least 75% reduction in NOS1AP relative to the non-targeting control.

For rescue experiments, sgRNA-resistant full-length human NOS1AP constructs encoding either wild-type NOS1AP or the CAPTOR-1-resistant Y16A/F42A variant were expressed from an EF1α-driven lentiviral vector. Guide resistance was introduced through synonymous substitutions within the sgRNA recognition sequence without altering the NOS1AP amino-acid sequence. Rescue constructs were introduced before NGN2 induction and expression was titrated to 80-120% of endogenous NOS1AP in non-targeting-control neurons, as measured by targeted proteomics. Empty-vector, wild-type-rescue, and Y16A/F42A-rescue cells underwent identical transduction, selection, and differentiation schedules. Experiments were performed using both NOS1AP-directed sgRNAs, and the reported effects were required to reproduce with each guide.

### Endogenous NOS1AP complexes

Neurons were treated with vehicle or **CAPTOR-1** at the indicated concentration for 2 h, washed twice with ice-cold phosphate-buffered saline, and lysed for 30 min at 4°C in 20 mM HEPES (pH 7.4), 150 mM NaCl, 0.2% NP-40 substitute, 1 mM EDTA, and protease and phosphatase inhibitor cocktails. Lysates were clarified by centrifugation at 16,000 × g for 15 min at 4°C, and protein concentrations were determined by bicinchoninic acid assay. For each immunoprecipitation, 500 µg of protein was adjusted to 500 µL and precleared with 20 µL of magnetic Protein G beads for 30 min at 4°C. Precleared lysates were incubated overnight at 4°C with 2 µg of mouse monoclonal anti-NOS1AP antibody (Santa Cruz Biotechnology, sc-374504), followed by 25 µL of magnetic

Protein G beads for 2 h at 4°C. Mouse IgG isotype and bead-only controls were processed in parallel. Beads were washed four times with lysis buffer and once with detergent-free buffer. Co-immunoprecipitated tau and captured NOS1AP were quantified by parallel-reaction-monitoring mass spectrometry using NOS1AP- and tau-specific proteotypic peptides and corresponding stable-isotope-labelled internal standards. Peak areas were integrated using Skyline, and peptide identity was confirmed by retention-time and fragment-ion matching to the heavy standards. Tau abundance was normalized first to immunoprecipitated NOS1AP and then to the vehicle condition within each donor and differentiation. The endogenous NOS1AP-nNOS complex was quantified from matched native lysates by AlphaLISA. Carrier-free rabbit monoclonal anti-NOS1AP antibody (clone EPR14519[B], Abcam, ab236134) was biotinylated using NHS-PEG4-biotin and detected with Alpha Streptavidin Donor Beads (Revvity, 6760002). Carrier-free rabbit monoclonal anti-nNOS antibody (clone EPR20372-53, Abcam, ab244742) was conjugated directly to AlphaLISA Unconjugated Acceptor Beads (Revvity, 6772001) according to the manufacturer’s protocol. Donor and acceptor beads were each used at 20 µg/mL. Lysate input was maintained within the experimentally established linear range. Three donor lines, each subjected to two independent neuronal differentiations, were analyzed. Technical replicates were averaged within each differentiation, and the donor-level mean was used as the statistical unit.

### Amyloid beta neuronal injury assay

Synthetic human amyloid-β_1-42_ peptide (AnaSpec, AS-20276) was initially dissolved in hexafluoroisopropanol (Sigma-Aldrich, 105228) to 1 mg/mL, aliquoted, and evaporated to dryness under a gentle nitrogen stream. Peptide films were stored at −80°C and used only once. Immediately before oligomer preparation, the peptide film was resuspended in anhydrous DMSO to 5 mM and diluted to 100 µM in phenol-red-free neuronal medium. The solution was incubated for 24 h at 4°C without agitation and then diluted directly into neuronal cultures to a final amyloid-β_1-42_ concentration of 5 µM. Vehicle controls received the corresponding concentration of DMSO.

Neurons were exposed for 24 h. **CAPTOR-1** (3 µM) was added 2 h before amyloid-β and remained present throughout the exposure period; the final DMSO concentration was matched across all conditions and did not exceed 0.1%. ATP-dependent viability was quantified using CellTiter-Glo 2.0 Luminescent Cell Viability Assay (Promega, G9241) according to the manufacturer’s instructions. Reagent was added at a 1:1 ratio to the culture medium, plates were mixed for 2 min and equilibrated for 10 min at room temperature, and luminescence was recorded using a Tecan Spark reader. Lactate dehydrogenase release was measured in conditioned medium using the CytoTox 96 Non-Radioactive Cytotoxicity Assay (Promega, G1780) and expressed relative to matched wells subjected to complete detergent-mediated lysis. Nitrite was quantified using the Griess Reagent System (Promega, G2930) against a sodium-nitrite standard curve and normalized to total cellular protein. Assays used three donor lines with two independent neuronal differentiations per donor. Technical wells were averaged within each differentiation, and the donor-level mean was used for statistical inference.

### Tau fibril preparation and neuronal seeding

Recombinant human tau-441 (2N4R) carrying the P301S substitution (StressMarq Biosciences, SPR-473) was diluted to 10 µM in aggregation buffer containing 100 mM sodium acetate (pH 7.0), 2 mM dithiothreitol, and unfractionated heparin sodium (Sigma-Aldrich, H3393) at a 4:1 tau-to-heparin mass ratio. Reactions were incubated for 72 h at 37°C with orbital agitation at 1,000 rpm in a thermomixer. Fibrils were collected by ultracentrifugation at 100,000 × *g* for 30 min at 4°C, washed twice with phosphate-buffered saline, and resuspended in sterile phosphate-buffered saline at 1 mg/mL. Immediately before neuronal exposure, fibrils were fragmented on ice by probe sonication at 20% amplitude using 30 cycles of 1 s on and 1 s off. Aliquots were prepared after sonication, stored at −80°C, and subjected to no more than one freeze-thaw cycle. For sedimentation analysis, matched aliquots were centrifuged at 100,000 × *g* for 30 min, and tau concentrations in the supernatant and pellet fractions were quantified by BCA assay. Amyloid formation was assessed by thioflavin-T fluorescence using 10 µM thioflavin-T, with excitation at 440 nm and emission recorded at 485 nm. PFF preparations were accepted only when at least 80% of tau partitioned into the pellet and the thioflavin-T signal was at least fivefold higher than that of the corresponding monomer preparation. Human NGN2-induced neurons were exposed to PFFs at 100 nM tau monomer equivalent for 72 h. **CAPTOR-1** at 0.3, 1, or 3 µM was added 2 h before PFF exposure and remained present throughout the experiment. The NOS1AP-CRISPRi plus compound condition received 3 µM **CAPTOR-1**. Vehicle and monomeric-tau controls were processed in parallel. Three independently prepared neuronal culture batches were analyzed, with two technical wells per condition; technical wells were averaged before batch-level statistical analysis.

### Tau aggregation and seeding assays

Cell-associated tau aggregates were quantified using the HTRF Tau Aggregation Kit (Revvity, 6FTAUPEG). Neurons were washed with ice-cold phosphate-buffered saline and lysed in the kit-compatible lysis buffer supplemented with protease and phosphatase inhibitors. Lysates were clarified at 16,000 × *g* for 15 min at 4°C and normalized by total protein concentration. Equal amounts of lysate were transferred to white low-volume 384-well plates, combined with the donor-and acceptor-labelled tau antibodies supplied with the kit, and incubated overnight at room temperature according to the manufacturer’s protocol. Time-resolved fluorescence was measured at 620 and 665 nm after excitation at 320 nm, and the HTRF ratio was calculated as 10^4^ × *F*_665_/*F*_620_ after subtraction of the reagent-blank signal. Total tau was quantified from matched lysates using the HTRF Total MAP-Tau Kit (Revvity, 63ADK008PEG). Aggregate signals were normalized to total protein, while total-tau measurements were used to confirm that treatment effects were not attributable to loss of tau protein. Tau-seeding activity was measured using Tau RD P301S FRET Biosensor cells (ATCC CRL-3275). Cells were maintained in DMEM containing 10% fetal bovine serum, 1% GlutaMAX, and 1% penicillin-streptomycin at 37°C and 5% CO_2_ and were confirmed to be mycoplasma-free. Biosensor cells were seeded at 35,000 cells per well in 96-well plates 24 h before treatment. Neuronal lysates were normalized to total protein, and 10 µg of protein was combined with 1.5 µL Lipofectamine 2000 (Thermo Fisher Scientific, 11668019) in Opti-MEM Reduced Serum Medium (Thermo Fisher Scientific, 31985070) for 20 min at room temperature before addition to the biosensor cells. After 48 h, cells were detached using TrypLE Express, washed, and analyzed by flow cytometry. CFP was detected after 405-nm excitation, YFP after 488-nm excitation, and FRET using 405-nm excitation with emission collected near 525 nm. Single cells were identified using forward- and side-scatter pulse geometry, and nonviable cells were excluded with a fixable viability dye. Spectral compensation was established using CFP-only and YFP-only controls, and the FRET-positive gate was fixed using unseeded cells exposed to Lipofectamine alone. Integrated FRET density was calculated as the percentage of FRET-positive cells multiplied by their median FRET intensity and normalized to the unseeded control. Two technical wells were averaged for each of three independently prepared neuronal culture batches.

### Commercial patient iPSC lines

The validation cohort comprised four AD patient-derived iPSC lines and four neurologically normal control iPSC lines obtained from the Coriell Institute. Each line represented an independent donor. Diagnosis and classification as sporadic AD were based on the clinical and genetic information provided by Coriell. The control and sporadic-AD groups were frequency-matched for sex (two females and two males per group) and age at biospecimen collection (controls, 70.8 ± 5.1 years; sporadic AD, 72.3 ± 4.6 years; Welch’s *P* = 0.69). All lines had supplier-confirmed identity, pluripotency, and karyotype and were confirmed to be mycoplasma-free before use. Each line underwent two independent NGN2 neuronal differentiations. Neurons were treated for 24 h with vehicle or **CAPTOR-1** at 1 or 3 µM. ATP-dependent viability, pTau217 abundance, tau-seeding activity, network-burst activity, and nitrite production were measured using cells or samples from the same differentiation batch. pTau217 was quantified by electrochemiluminescent immunoassay and normalized to total protein. Network activity was measured using multielectrode arrays and reported as the mean network-burst rate during a prespecified stable 10-min recording period at 37°C, normalized to the mean of vehicle-treated control lines. Technical replicates were averaged within each differentiation, and the donor-line mean across the two differentiations was used as the biological replicate.

### Phosphoproteomics

Phosphoproteomic profiling was performed using four neurologically normal iPSC donor lines (from the Coriell Institute), with two independent NGN2 neuronal differentiations per line. For each donor, the experimental conditions were unseeded control neurons, neurons exposed to P301S tau PFFs, PFF-exposed neurons treated with 3 µM **CAPTOR-1** for 24 h, PFF-exposed neurons subjected to NOS1AP CRISPRi, and PFF-exposed NOS1AP-CRISPRi neurons rescued with the **CAPTOR-1**-resistant Y16A/F42A NOS1AP variant and treated with 3 µM **CAPTOR-1** for 24 h. Cells were washed twice with ice-cold phosphate-buffered saline and lysed in 8 M urea, 50 mM triethylammonium bicarbonate (pH 8.5), 1 mM sodium orthovanadate, 10 mM sodium fluoride, 10 mM β-glycerophosphate, and EDTA-free protease inhibitor cocktail. Lysates were disrupted by brief probe sonication on ice and clarified at 16,000 × *g* for 15 min at 4°C. Protein concentration was determined by BCA assay. For each sample, 100 µg of protein was reduced with 5 mM dithiothreitol for 30 min at 37°C and alkylated with 15 mM iodoacetamide for 30 min at room temperature in the dark. Excess iodoacetamide was quenched with 5 mM dithiothreitol. Proteins were digested with Lys-C (FUJIFILM Wako, 125-05061) at an enzyme-to-protein ratio of 1:100 for 3 h at 37°C. Samples were then diluted fourfold with 50 mM triethylammonium bicarbonate and digested overnight with sequencing-grade modified trypsin (Promega, V5111) at a 1:50 enzyme-to-protein ratio. Digestion was stopped with 1% formic acid, and peptides were desalted using C18 solid-phase extraction cartridges. Equal peptide amounts were labeled with TMTpro 18plex reagents (Thermo Fisher Scientific, A52045), pooled, and fractionated. Phosphopeptides were enriched with Fe-NTA resin and analyzed by nanoLC-MS/MS on an Orbitrap Exploris 480. Spectra were searched against the reviewed human UniProt database with a 1% peptide and protein false-discovery rate. Reporter-ion intensities were corrected for isotopic impurities, normalized to total peptide abundance, and summarized at the phosphosite level. Tau T217, T231, and S202 formed the prespecified primary family.

### In vitro ADME and safety assays

**CAPTOR-1** kinetic solubility was measured after 2 h in phosphate buffer pH 7.4 at a nominal compound concentration of 100 µM, with 1% DMSO, at 25°C, by filtration followed by LC-MS/MS. Intrinsic clearance was measured in mouse and human liver microsomes (from Sigma Aldrich) at 0.5 mg mL^−1^ protein and 1 µM compound in the presence of an NADPH-regenerating system; disappearance was sampled at 0, 5, 10, 20, 40, and 60 min and fit to a first-order model. Parallel incubations without NADPH and reference-compound incubations with verapamil were included as controls. Microsomal half-life and intrinsic clearance were calculated from the depletion-rate constant. Passive blood-brain barrier permeability was measured with a PAMPA-BBB assay using porcine brain lipid at pH 7.4 after a 4-h incubation; caffeine and corticosterone served as low- and high-permeability controls, respectively. hERG channel inhibition was measured at 10 µM by automated whole-cell patch clamp in a stable hERG-expressing HEK293 cell line at physiological temperature; cisapride was the positive control. Each ADME assay was performed in at least three independent incubations with duplicate analytical measurements, with appropriate positive controls.

### Mouse pharmacokinetics

Mice were housed and handled according to the guidelines approved by the Institutional Animal Care and Use Committee (IACUC) of our institution (protocol 2024-0010). Pharmacokinetics were evaluated in adult C57BL/6J mice (The Jackson Laboratory, strain no. 000664; 10-12 weeks old) after a single 30 mg kg^−1^ oral dose of **CAPTOR-1** formulated in 0.5% methylcellulose and 0.1% Tween-80 in purified water and administered at 10 mL kg^−1^. Four animals were sampled at each time point, two females and two males. Blood and perfused whole brain were collected at 0.25, 0.5, 1.5, 4, 8, and 24 h. Animals were terminally sampled at one time point, and brains were collected after transcardial perfusion with ice-cold phosphate-buffered saline to minimize residual intravascular drug. Plasma was separated within 30 min; brain was weighed and homogenized in four volumes of ice-cold water. **CAPTOR-1** was quantified by a validated LC-MS/MS method with a stable-isotope internal standard and matrix-matched calibration covering 1-5,000 ng mL^−1^ for plasma and 0.5-2,500 ng g^−1^ for brain; accuracy and precision were required to remain within ±15% except at the lower limit of quantification, where ±20% was accepted. Plasma and brain concentration-time profiles were constructed from the mean concentration at each terminal sampling time. Noncompartmental analysis was performed using Phoenix WinNonlin, version 8.4. AUC_0–24_ was calculated for each matrix from the composite mean profile using the linear-up/log-down trapezoidal method. Brain partitioning was expressed as Kp,AUC = _AUCbrain,0–24_/AUC_plasma,0–24_. Because different animals contributed to each time point, Kp,AUC represents a single composite estimate without an inter-animal standard deviation. Brain-to-plasma concentration ratios at individual time points were calculated from matched brain and plasma concentrations within each animal and are presented as mean ± S.D. A separate cohort received **CAPTOR-1** intravenously at 2 mg kg^−1^ in the same formulation. Plasma **CAPTOR-1** was quantified over 24 h using the same LC-MS/MS method. Absolute oral bioavailability (%) = [(oral AUC ÷ oral dose) ÷ (intravenous AUC ÷ intravenous dose)] × 100.

### Brain target engagement

C57BL/6J mice, 10-12 weeks old and balanced by sex, received vehicle or a single oral dose of **CAPTOR-1** at 3, 10, or 30 mg kg^−1^ (n = 6 per dose; three females and three males). Animals were euthanized 2 h after dosing. Cortex and hippocampus were rapidly dissected, combined within animals, and homogenized under native conditions in 20 mM HEPES, pH 7.4, 150 mM NaCl, 0.2% NP-40 substitute, and protease and phosphatase inhibitors. For CETSA, aliquots were heated for 3 min at the empirically selected aggregation temperature of 52°C, cooled for 3 min at room temperature, and centrifuged at 20,000 × g for 20 min at 4°C. Soluble NOS1AP in the supernatant was quantified by a sandwich AlphaLISA assay and expressed relative to the vehicle mean. Recombinant NOS1AP standards were included on each plate, and samples were analyzed within the assay’s established linear range. Matched unheated aliquots confirmed equal total NOS1AP input. NOS1AP-tau and NOS1AP-nNOS complexes were quantified in unheated homogenates by AlphaLISA. For each assay, homogenate containing 10 µg total protein was incubated with a biotinylated anti-NOS1AP antibody (1 nM) and either an anti-tau or anti-nNOS antibody (1 nM) for 60 min at room temperature. AlphaLISA acceptor beads directly conjugated to the anti-tau or anti-nNOS antibody were added to a final concentration of 20 µg mL^−1^ and incubated for 30 min at room temperature. Streptavidin donor beads were then added to 20 µg mL^−1^, followed by incubation for 60 min at room temperature in the dark. Luminescence was measured at 615 nm after 680-nm excitation using an EnVision multimode plate reader. Bead-only, single-antibody, isotype-control, and excess-antigen competition controls were included. Signals were background-subtracted, confirmed to fall within the linear protein-input range, and normalized to total NOS1AP measured in the corresponding unheated homogenate.

### Animals and welfare

Mice were housed and handled according to the guidelines approved by the Institutional Animal Care and Use Committee (IACUC) of our institution (protocol 2024-0010). PS19 mice [The Jackson Laboratory stock 008169] and APP/PS1 mice [The Jackson Laboratory, strain no. 005864] were maintained with wild-type littermates under a 12-h light-dark cycle at 22 ± 2°C and 40-60% relative humidity, with food and water available ad libitum. Genotypes were confirmed by supplier-recommended PCR assays before enrollment. Both sexes were used. Each efficacy study contained four groups of 16 animals: wild-type vehicle, wild-type **CAPTOR-1**, transgenic vehicle, and transgenic **CAPTOR-1**, with eight females and eight males per group. Investigators remained blinded to genotype and treatment allocation during behavioral testing, sample processing, data acquisition, and analysis; group identities were revealed only after the prespecified analyses were completed.

### Chronic efficacy dosing

PS19 mice received **CAPTOR-1** at 30 mg kg^−1^ or vehicle by oral gavage once daily from 6 to 8 months of age. APP/PS1 mice received the same regimen from 8 to 10 months of age. **CAPTOR-1** was formulated in 0.5% methylcellulose and 0.1% Tween-80 in purified water and administered at 10 mL kg^−1^; vehicle groups received an identical volume of formulation without compound. Formulations were prepared weekly, stored at 4°C protected from light, and verified for concentration and stability by LC-MS/MS. Dose volume was adjusted weekly to body weight. Animals were monitored daily for clinical condition and at least twice weekly for body weight. Food consumption and clinical observations were recorded throughout treatment. Behavioral testing occurred during the final two treatment weeks, with dosing performed after testing on behavioral days to prevent acute exposure or handling from influencing performance. Terminal tissue and cerebrospinal fluid collection occurred 2 h after the final dose. Terminal plasma and brain **CAPTOR-1** concentrations were determined by LC-MS/MS as exposure-confirmation endpoints.

### Contextual fear conditioning

Mice were habituated to the testing room for at least 30 min. During training, each animal explored the conditioning chamber for 2 min before receiving three 0.7-mA foot shocks of 2 s duration separated by 60 s. Animals remained in the chamber for 60 s after the final shock. Twenty-four hours later, mice were returned to the same chamber for a 5-min context test without shock.

Freezing was scored automatically from activity traces using one prespecified threshold validated by manual scoring of a randomly selected, blinded subset before analysis and expressed as percentage of the session. Freezing was defined as the absence of movement other than respiration for at least 1 s. Chambers were cleaned between animals with 70% ethanol, and test order was balanced across genotype, treatment, and sex. Training-session activity and shock reactivity were examined as controls for baseline locomotor or sensory differences.

### Radial arm water maze

Spatial learning in APP/PS1 mice was evaluated in a six-arm radial-arm water maze containing opaque water maintained at 22 ± 1°C, with a submerged escape platform positioned 1 cm below the surface and fixed in one goal arm. Distinct distal visual cues were maintained in constant positions throughout testing. Mice completed five acquisition trials per day for two consecutive days, with randomized start arms and a maximum trial duration of 60 s. Mice failing to locate the platform within 60 s were guided to it and allowed to remain there for 15 s; intertrial intervals were 10-15 min. An incorrect arm entry was counted when the mouse entered a non-goal arm with all four paws. Errors from the five Day 2 trials were averaged for each animal. Swim speed and escape latency were recorded as secondary measures to identify motor or motivational confounding. Experimenters conducting and scoring the task were blinded to group.

### Hippocampal electrophysiology

Acute transverse hippocampal slices, 350 micrometers thick, were prepared in ice-cold oxygenated sucrose cutting solution containing sucrose 206 mM, KCl 2.8 mM, CaCl_2_ 1 mM, MgCl_2_, 2 mM, NaH_2_PO_4_1.25 mM, NaHCO_3_ 26 mM, and glucose 10 mM, continuously equilibrated with 95% O_2_/5% CO_2_, and recovered for at least 60 min in artificial cerebrospinal fluid at 32°C. The recording solution contained NaCl 124 mM, KCl 2.8 mM, CaCl_2_ 2 mM, MgSO_4_ 1 mM, NaH_2_PO_4_ 1.25 mM, NaHCO_3_ 26 mM, and glucose 10 mM and was perfused at 2-3 mL min^−1^. Field excitatory postsynaptic potentials were recorded from stratum radiatum of CA1 while Schaffer collaterals were stimulated at 0.05 Hz. Baseline responses were adjusted to 40-50% of the maximal slope and recorded for 20 min. Slices with baseline drift exceeding 10% or unstable fiber-volley amplitudes were excluded according to prespecified criteria. Long-term potentiation was induced by theta-burst stimulation consisting of four trains separated by 20 s; each train contained ten bursts at 5 Hz, with four pulses at 100 Hz per burst. Responses were recorded for 60 min after induction. The primary endpoint was the mean fEPSP slope during minutes 50–60, expressed as percentage of the pre-induction baseline. One prespecified technically acceptable slice per animal was used for inference. Slice selection and electrophysiological analysis were completed while investigators remained blinded to genotype, treatment, and sex.

### Mouse biochemical endpoints

For PS19 mice, detergent-soluble hippocampal homogenates prepared in 20 mM HEPES, pH 7.4, 150 mM NaCl, 1% Triton X-100, and protease and phosphatase inhibitors were analyzed for tau aggregation by HTRF, pTau181 and pTau231 by electrochemiluminescent immunoassay, and tau-seeding activity with the flow-cytometric FRET biosensor assay. Phospho-tau concentrations were normalized to total tau and total homogenate protein. Tau-seeding measurements were performed using equal amounts of total protein, and integrated FRET density was calculated as the percentage of FRET-positive cells multiplied by their median FRET intensity. For APP/PS1 mice, guanidine-extractable cortical Aβ_42_ was measured by sandwich ELISA and normalized to total tissue protein; cortical nitrite was measured by the Griess assay. Aβ_42_ extraction used 5 M guanidine hydrochloride in 50 mM Tris-HCl, pH 8.0, followed by centrifugation at 20,000 × g for 20 min at 4°C. Cerebrospinal fluid neurofilament light was measured in both models by ultrasensitive single-molecule immunoassay. Samples were randomized across assay plates, and each plate contained a pooled quality-control sample, blank, calibration curve, and duplicate technical measurements. Samples exceeding the assay range were diluted and reanalyzed.

Samples were assigned coded identifiers, and each animal contributed one value per endpoint after technical replicates were averaged. Assay operators remained blinded until quality control and prespecified data exclusions were finalized.

### Statistical analysis

Statistical analyses were performed in GraphPad Prism 10 and R 4.3. Continuous data are shown as individual biological replicates with mean and standard deviation unless stated otherwise. Binding curves were fit by nonlinear regression; affinities were analyzed after log10 transformation when compared across variants. Two-group donor or vendor comparisons used paired t tests when measurements were naturally paired and Welch t tests otherwise. Comparisons of multiple proteins with NOS1AP used Dunnett correction. Concentration series and prespecified endpoint families used Holm adjustment as identified in the figure legends. Patient-neuron analyses used donor-level means of the two differentiations. Mouse endpoints were analyzed with genotype, treatment, sex, and the genotype-by-treatment interaction in a linear model, with breeding cohort included as a blocking factor; planned within-genotype contrasts were Holm-adjusted across endpoints. Sex-by-treatment interactions were examined as secondary analyses but were not used to define efficacy. All tests were two sided. *P* less than 0.05 after the specified multiplicity correction was considered significant. No technical replicate was treated as an independent biological replicate.

## Author contributions

The manuscript was written through contributions of all authors. All authors have given approval to the final version of the manuscript. Conceptualization: M.T.G. Methodology: A.N.A., S.C., and Y.G. Investigation: A.N.A., S.C., and Y.G. Formal analysis: A.N.A., S.C., and Y.G. Visualization: A.N.A., S.C., and Y.G. Supervision: M.T.G. and H.D. Writing-original draft: A.N.A., S.C., Y.G., M.T.G., and H.D. Writing-review and editing: M.T.G. Funding acquisition: M.T.G. and H.D. Data curation: A.N.A., S.C., and Y.G. Validation: A.N.A., S.C., and Y.G. Project administration: M.T.G.

## Data and materials availability

All data needed to evaluate the conclusions in the paper are present in the paper and/or the Supplementary Materials.

