## Supporting Information for "A First-in-Class, Brain-Penetrant Small Molecule Inhibitor of the NOS1AP-Tau Interaction Rescues Tau Pathology In Vivo and Improves Cognition"

*Electronic Supplementary Information for*

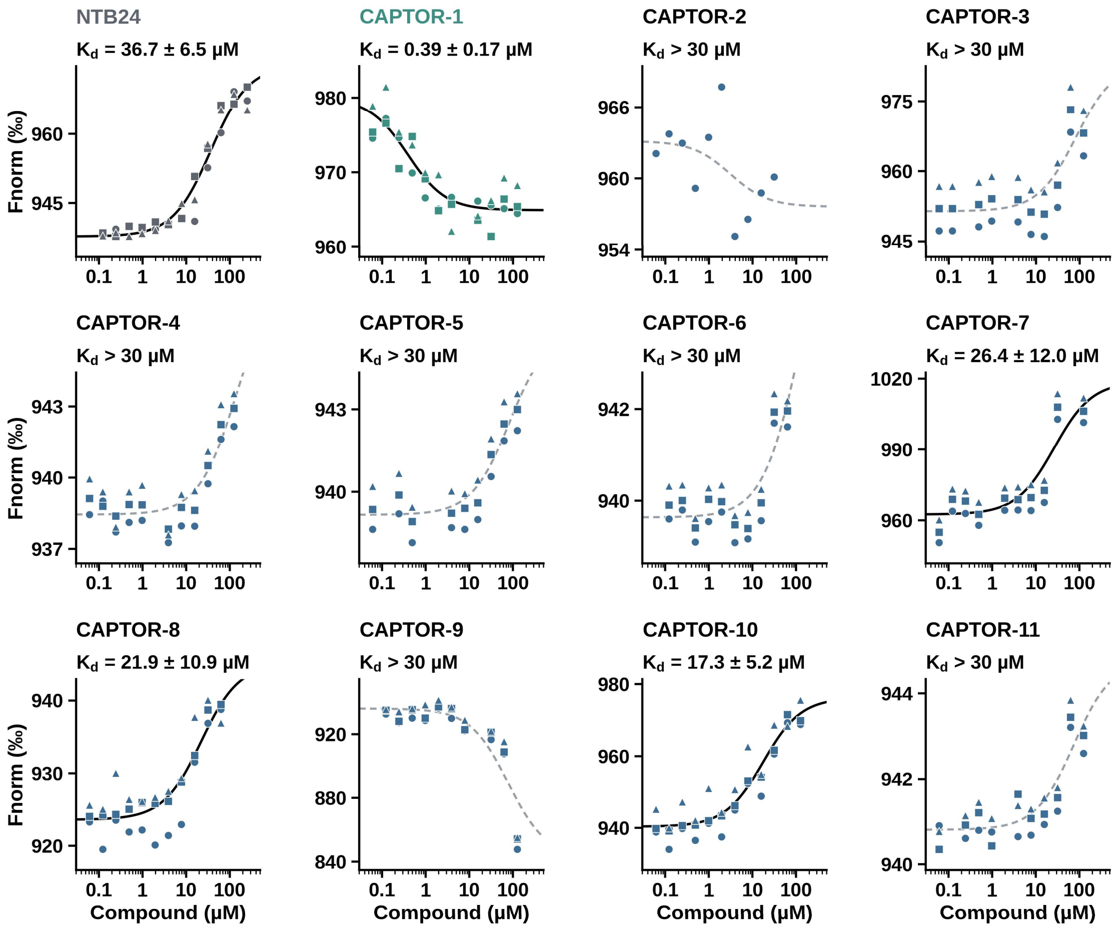


**Supplementary Figure 1. Raw MST data for the dose-dependent NOS1AP binding for all compounds.** Fnorm (‰) versus compound concentration for **NTB24** and the eleven analogs. All technical replicates are shown (n = 3, three symbols). Solid lines are the 1:1 binding fits used for the reported Kd (fitted value ± S.E. above each panel). Dashed lines are poorly constrained fits that were not used; for these compounds the affinity is reported as Kd > 30 µM. **CAPTOR-2** was tested only up to 31 µM. **CAPTOR-9** shows a large, non-saturating decrease in signal at ≥ 30 µM, which we interpret as nonspecific (e.g. aggregation or quenching) and not as binding.

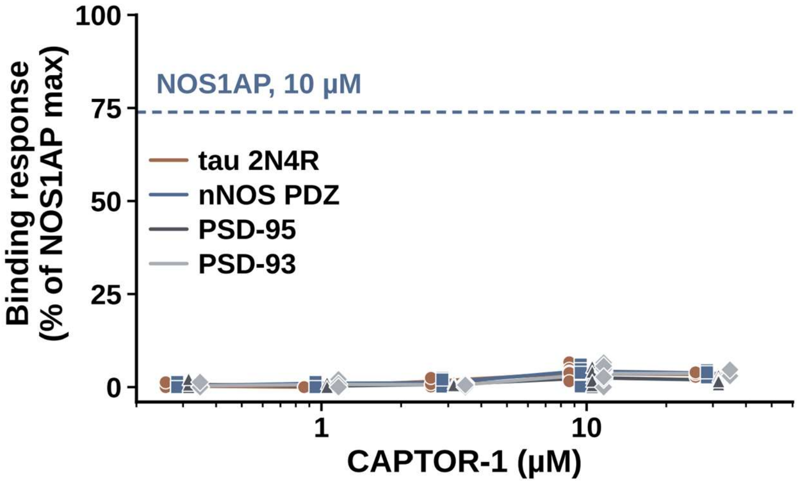


**Supplementary Figure 2. Concentration series for the CAPTOR-1 binding against four proteins.** Binding response of **CAPTOR-1** (0.3-30 µM) to tau 2N4R, nNOS PDZ, PSD-95 and PSD-93, as % of the NOS1AP maximum response. Individual replicates are shown (n = 4 per concentration); lines connect means. The dashed line is the NOS1AP response at 10 µM (73.9%). Mean responses remained below 5% at every concentration, with no concentration dependence.

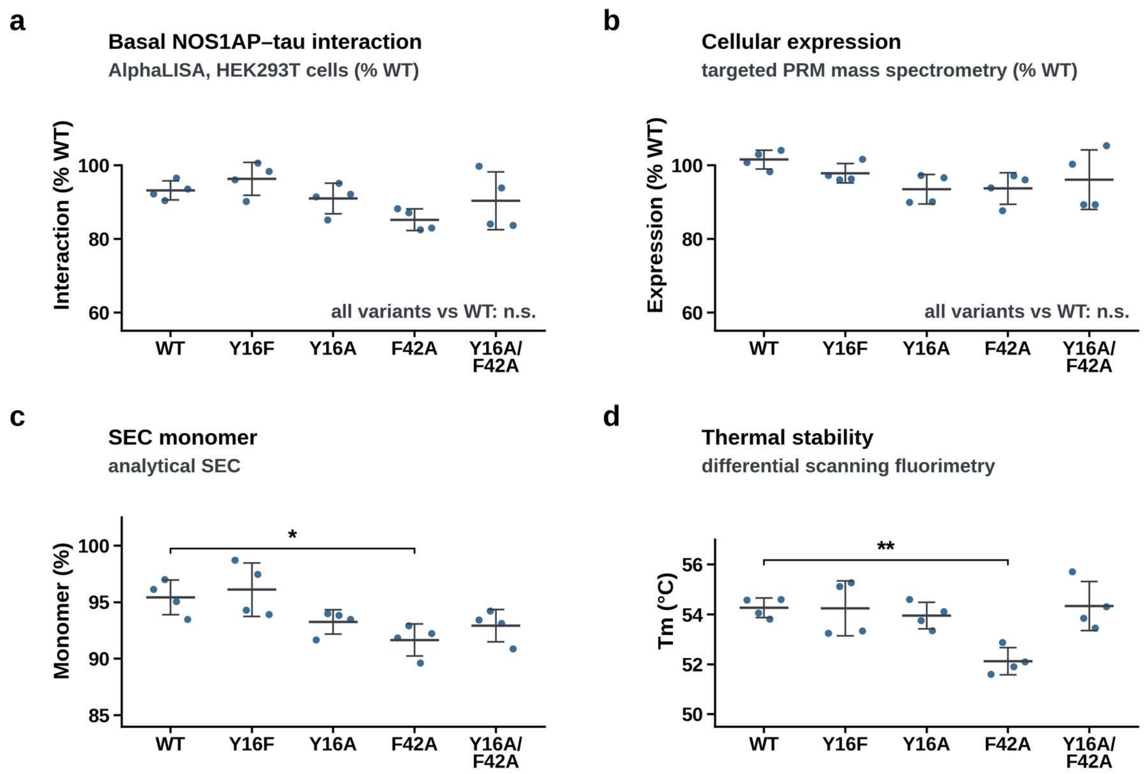


**Supplementary Figure 3. Integrity of the NOS1AP variants. a)** Basal NOS1AP-tau interaction (AlphaLISA, transfected HEK293T cells; % wild type). **b)** Cellular expression (targeted PRM mass spectrometry; % wild type). **c)** SEC monomer (analytical size-exclusion chromatography). **d)** Melting temperature (differential scanning fluorimetry). Points are independent protein preparations or transfections (n = 4); bars are mean ± S.D.; brackets show Dunnett's test versus wild type (* $P < 0.05$, ** $P < 0.01$); all other comparisons are not significant.

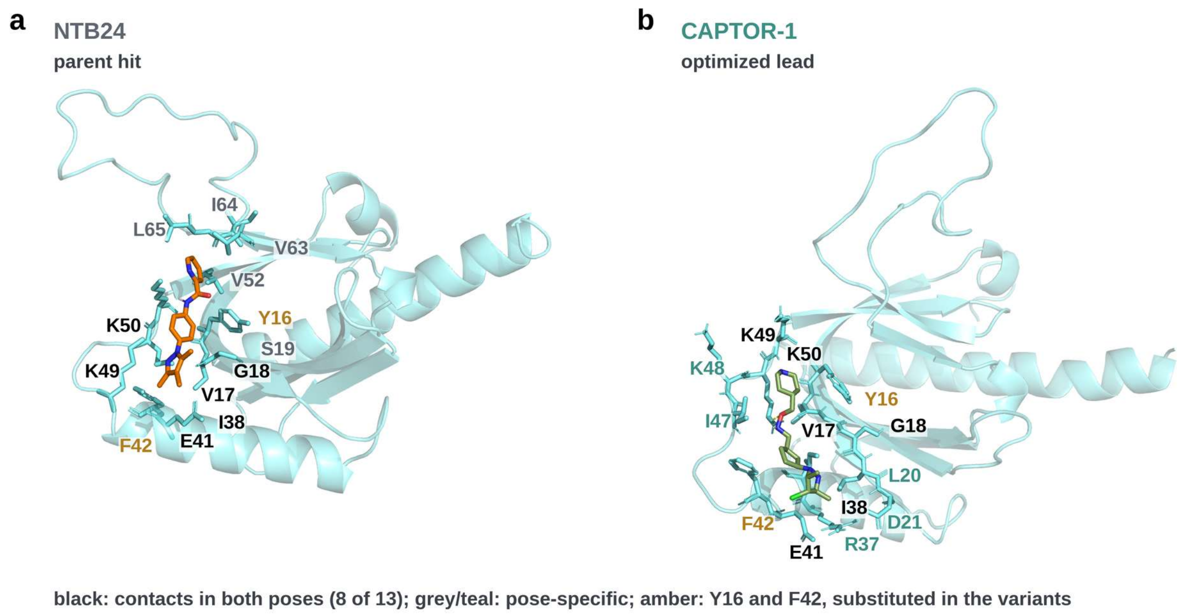


**Supplementary Figure 4. Docked and simulated poses of NTB24 and CAPTOR-1 to the NOS1AP homology model. a)** Pose of **NTB24** in the NOS1AP homology model (MD average structure, 80-100 ns). **b)** Pose of **CAPTOR-1** in the same model (MD average structure, 80-100 ns). Views are not superimposed. Labeled residues contact the ligand; eight of 13 contacts are shared (black); amber marks Y16 and F42.

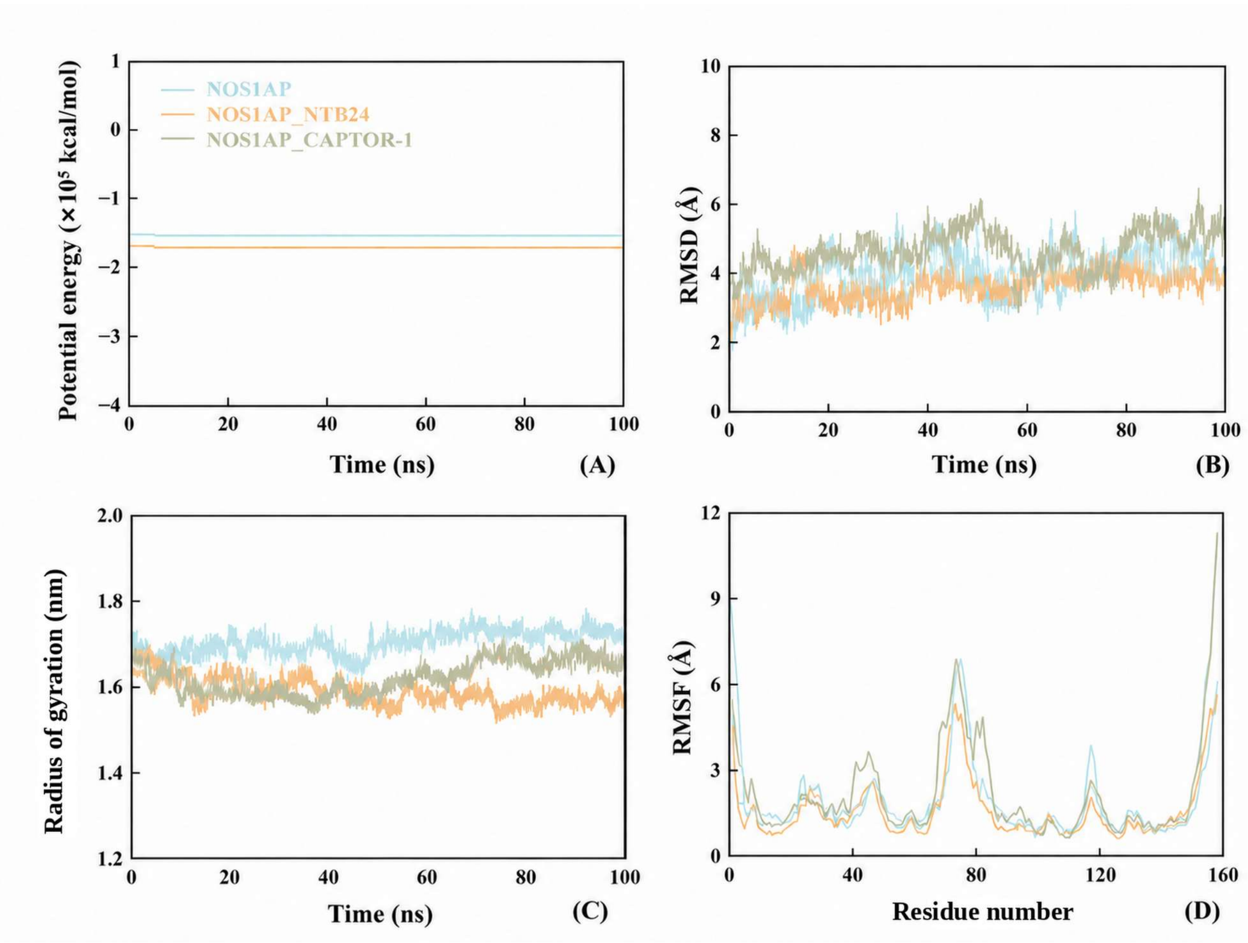


**Supplementary Figure 5. Stability of the simulated NOS1AP-NTB24 and NOS1AP-CAPTOR-1 complexes. a)** Potential energy. **b)** Backbone RMSD. **c)** Radius of gyration. **d)** Per-residue RMSF. Molecular dynamics of apo NOS1AP, NOS1AP with **NTB24** and NOS1AP with **CAPTOR-1** over 100 ns (one trajectory per system). Fluctuations are similar across the three systems, with the largest RMSF peaks in the same loop region (residues about 70-80) and at the termini.

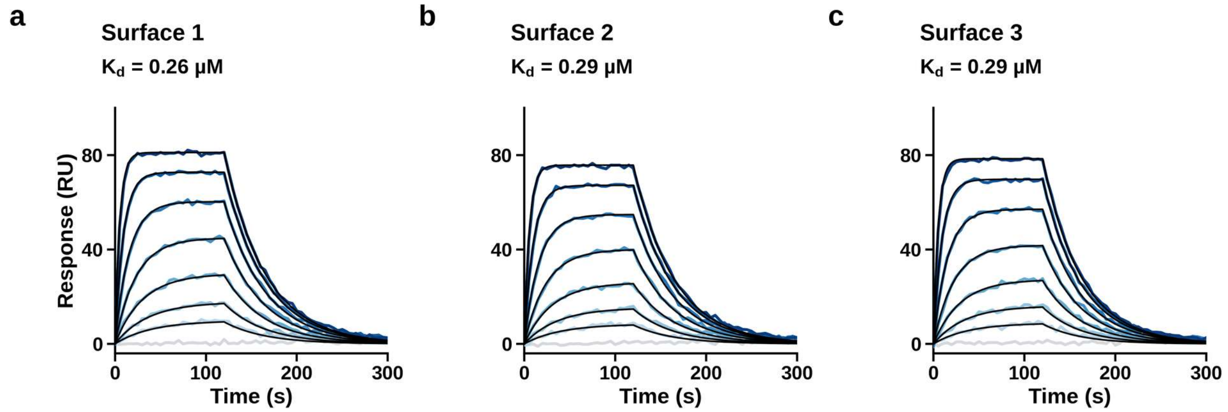


**Supplementary Figure 6. SPR sensorgrams for the three independent surfaces for the NOS1AP-tau interaction. a-c)** Surfaces 1-3 with global 1:1 kinetic fits (black). NOS1AP concentrations are 0.03-2 μM as in Fig. 4a (light grey, buffer injection); Kd per surface is from the kinetic fit.

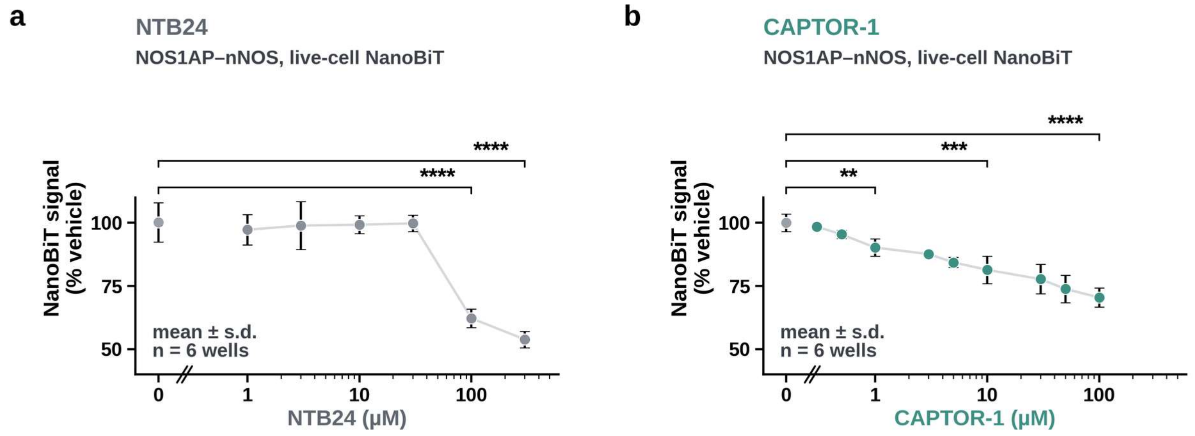


**Supplementary Figure 7. NanoBiT NOS1AP-nNOS counter-screen. a) NTB24** in live cells. **b) CAPTOR-1** in live cells. Mean ± S.D. of n = 6 (technical replicates) from one experiment; vehicle at left. Brackets: two-sided t-tests versus vehicle computed from the summary statistics, Holm-adjusted across concentrations (** $P < 0.01$, *** $P < 0.001$, **** $P < 0.0001$).

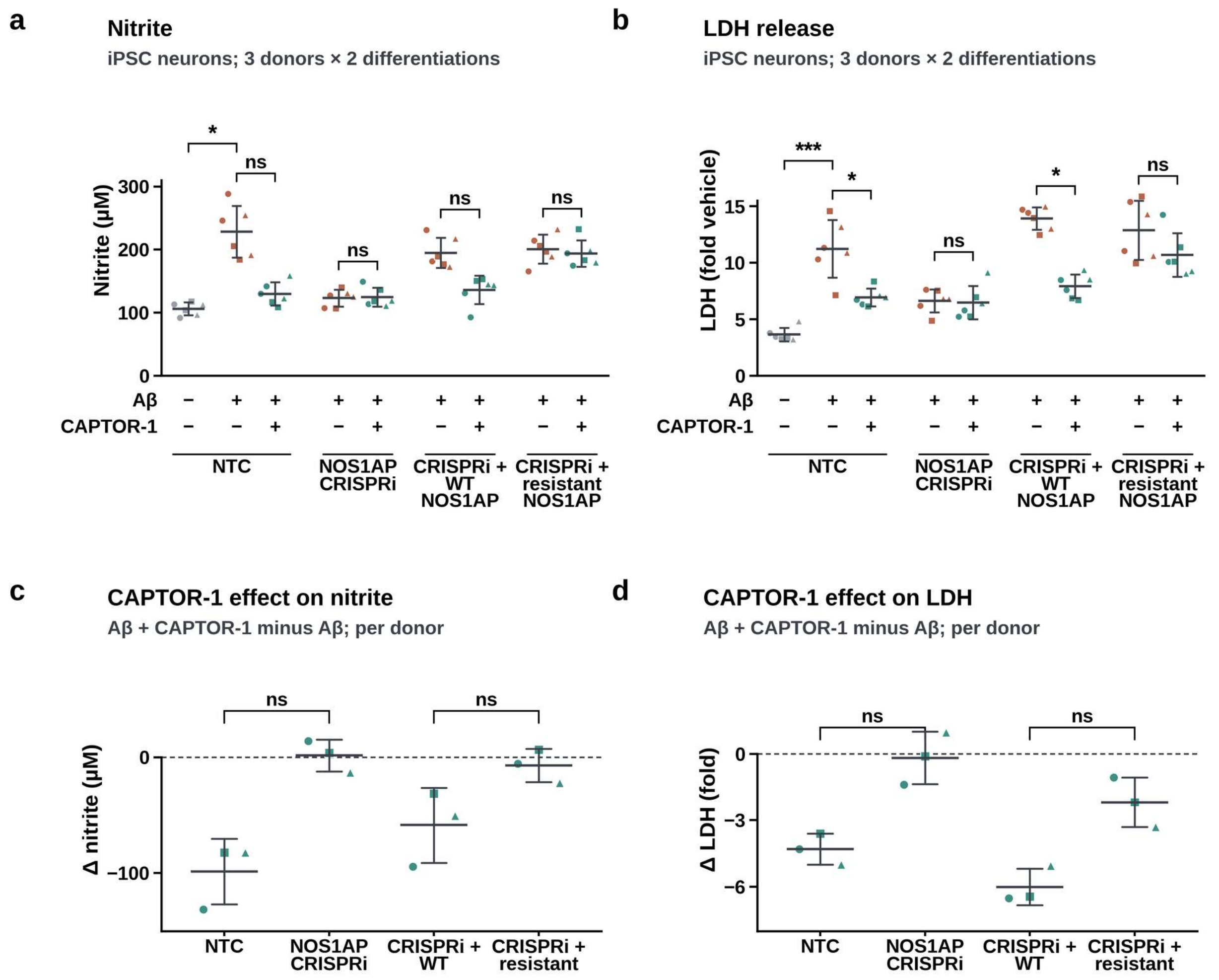


**Supplementary Figure 8. Nitrite and LDH release in the genotype-epistasis in vitro evaluation of CAPTOR-1. a)** Nitrite in iPSC-derived neurons exposed to Aβ, with non-targeting control (NTC) or NOS1AP CRISPRi, and CRISPRi neurons re-expressing wild-type or drug-resistant (Y16A/F42A) NOS1AP, with or without **CAPTOR-1** (design matrix below the axis). **b)** LDH release (fold vehicle) under the same conditions. **c) CAPTOR-1** effect on nitrite (Aβ + **CAPTOR-1** minus Aβ) per donor. **d) CAPTOR-1** effect on LDH release per donor. Points are observations (3 donors × 2 differentiations; symbols mark donors); bars are mean ± S.D. Brackets in **a** and **b**: paired t-tests on donor means (n = 3), Aβ effect unadjusted and **CAPTOR-1** effects Holm-adjusted across four genotypes (* $P < 0.05$, *** $P < 0.001$, ns not significant). Brackets in **c** and **d**: paired t-test between genotypes, Holm-adjusted across two contrasts.

a

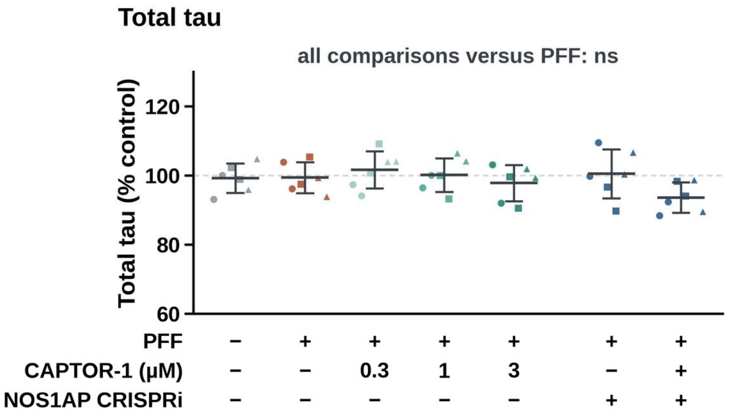


**Supplementary Figure 9. Total tau in the PFF seeding experiment. a)** Total tau (% of control) in neurons treated as in Fig. 6a,b (design matrix below the axis). Points are observations (3 culture batches × 2 replicates; symbols mark batches); bars are mean ± S.D. No comparison with PFF alone was significant (paired t-tests on batch means, Holm-adjusted).

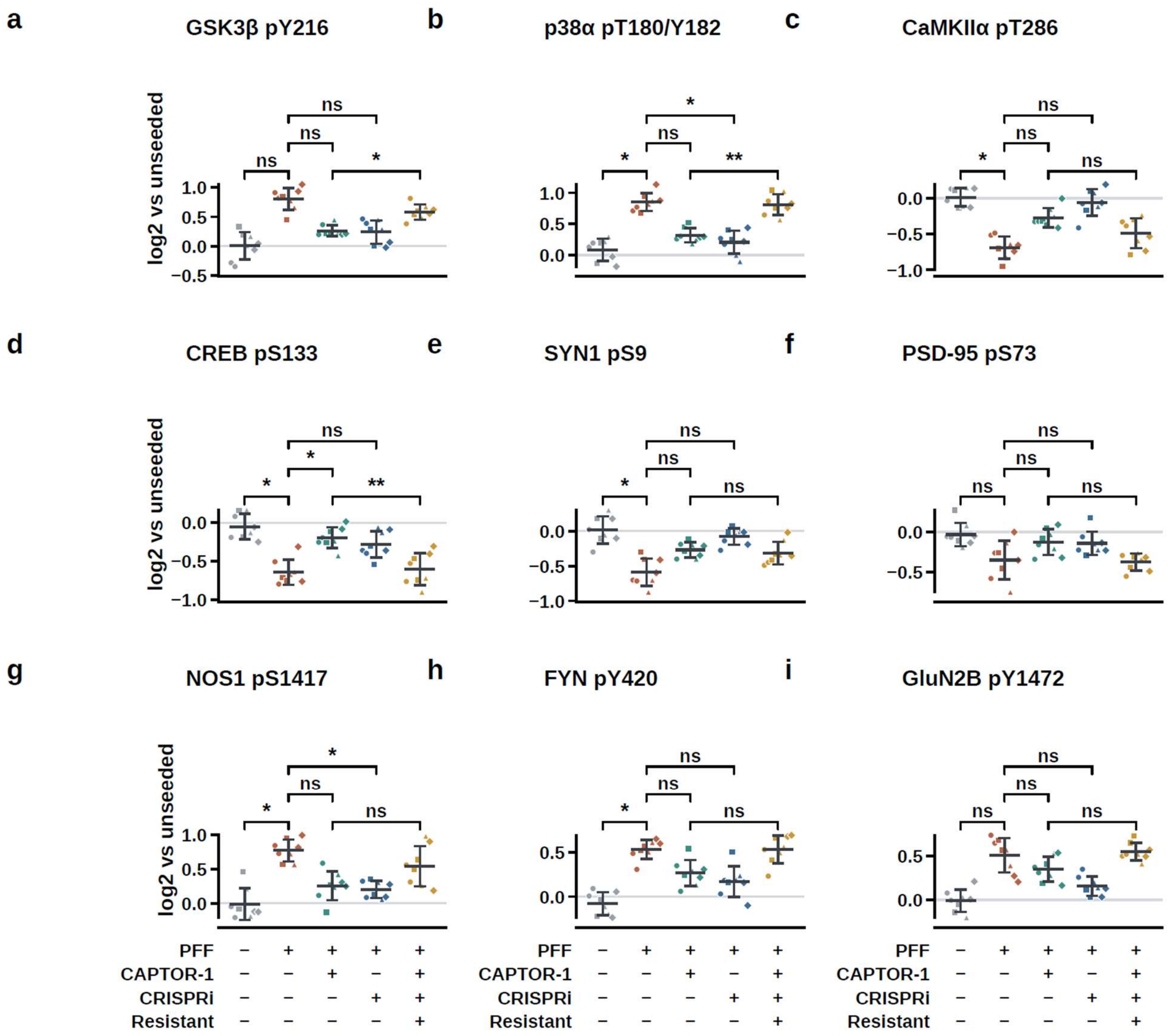


**Supplementary Figure 10. Phosphorylation of nine signaling sites in PFF-exposed neurons. a-i)** Phosphosite abundance (log2 versus control) for GSK3β pY216 **(a)**, p38α pT180/Y182 **(b)**, CaMKIIα pT286 **(c)**, CREB pS133 **(d)**, SYN1 pS9 **(e)**, PSD-95 pS73 **(f)**, NOS1 pS1417 **(g)**, FYN pY420 **(h)** and GluN2B pY1472 **(i)** in control, PFF-exposed neurons, PFF-exposed neurons + **CAPTOR-1**, PFF-exposed neurons + NOS1AP CRISPRi, and PFF-exposed neurons + CRISPRi + resistant NOS1AP (Y16A/F42A) + **CAPTOR-1** neurons (design matrix below the bottom row). Points are observations (4 donors × 2 differentiations; symbols mark donors); bars are mean ± S.D.; brackets, paired t-tests on donor means (n = 4), Holm-adjusted across the nine sites (exploratory family) (* $P < 0.05$, ** $P < 0.01$; ns, not significant).

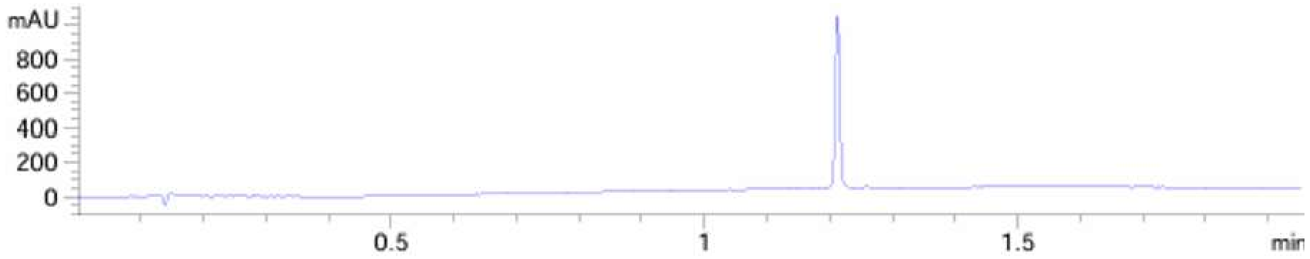


**Supplementary Figure 11.** HPLC trace of **CAPTOR-1**. Purity: >99% (UV detection at 254 nm). Analysis was performed on a C18 reversed-phase column using a water/acetonitrile gradient containing 0.1% formic acid, at a flow rate of 1.0 mL/min.

**Supplementary Table 1. Structures and MST binding affinities of NTB24 and its analogs for NOS1AP.** Structural change relative to **NTB24** (added or substituted groups highlighted in yellow; deletions not marked), molecular weight, MST Kd (fitted value ± S.E. of a 1:1 binding fit to three technical replicates per concentration), and highest concentration tested. Curve unsaturated or poorly constrained are reported as Kd> 30 µM.

| Compound | Structure | Structural change relative to NTB24 | MW (Da) | MST Kd (µM) | Max. conc. tested (µM) |
|---|---|---|---|---|---|
| **NTB24** | 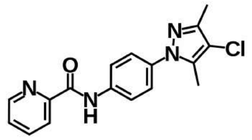 | Parent hit; 2-pyridyl amide (nitrogen ortho to carbonyl) | 326.8 | 36.7 ± 6.5 | 250.0 |
| **CAPTOR-1** | 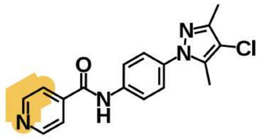 | 4-Pyridyl amide (nitrogen para) | 326.8 | 0.39 ± 0.17 | 125.0 |
| **CAPTOR-2** | 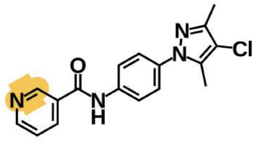 | 3-Pyridyl amide (nitrogen meta) | 326.8 | > 30 ‡ | 31.3 |
| **CAPTOR-3** | 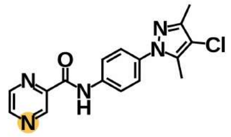 | Pyrazinyl amide (two ring nitrogens) | 327.8 | > 30 ‡ | 125.0 |
| **CAPTOR-4** | 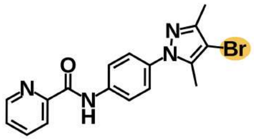 | Pyrazole 4-bromo in place of 4-chloro | 371.2 | > 30 ‡ | 125.0 |
| **CAPTOR-5** | 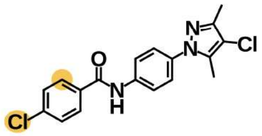 | 4-Chlorophenyl amide (no ring nitrogen) | 360.2 | > 30 ‡ | 125.0 |
| **CAPTOR-6** | 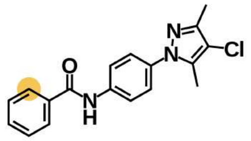 | Phenyl amide (no ring nitrogen) | 325.8 | > 30 ‡ | 62.5 |
| **CAPTOR-7** | 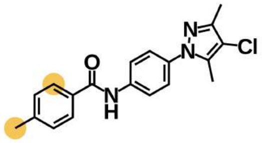 | 4-Methylphenyl amide (no ring nitrogen) | 339.8 | 26.4 ± 12.0 † | 125.0 |

| Compound | Structure | Structural change relative to NTB24 | MW (Da) | MST Kd (µM) | Max. conc. tested (µM) |
|---|---|---|---|---|---|
| **CAPTOR-8** | 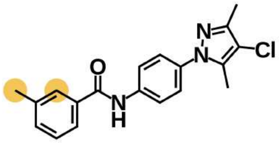 | 3-Methylphenyl amide (no ring nitrogen) | 339.8 | 21.9 ± 10.9 † | 62.5 |
| **CAPTOR-9** | 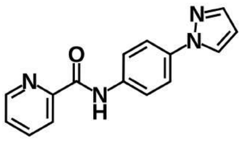 | Substituted-pyrazole core absent | 264.3 | > 30 ‡ | 125.0 |
| **CAPTOR-10** | 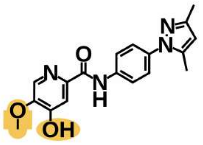 | Hydroxy-methoxypyridine amide; altered pyrazole substitution | 338.4 | 17.3 ± 5.2 | 125.0 |
| **CAPTOR-11** | 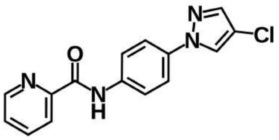 | Pyrazole lacking both methyl groups | 298.7 | > 30 ‡ | 125.0 |

Kd values were obtained by microscale thermophoresis (MST) using labeled NOS1AP. Data (three technical replicates per concentration) were fitted to a 1:1 binding model with free baseline and plateau by non-linear least squares; values are the fitted Kd ± standard error of the fit.
Added or substituted groups relative to NTB24 are highlighted in amber; deletions are not marked.
† Standard error of the fit is approximately 45–50% of the fitted value; the value should be regarded as approximate.
‡ The binding curve was unsaturated or poorly constrained over the concentration range tested, so a Kd could not be reliably determined; the affinity is reported as > 30 µM.

### Supplementary Table 2. CAPTOR-1-induced changes in deuterium uptake of NOS1AP

| Peptide | Residues | Region | ΔD uptake at 30 s (%) | ΔD uptake at 300 s (%) | ΔD uptake at 3,000 s (%) |
|---|---|---|---|---|---|
| P1 | 9-27 | N-terminal region | −2.42 ± 0.22 | −3.21 ± 0.06 | −3.99 ± 0.14 |
| P2 | 35-53 | N-terminal region | −1.38 ± 0.27 | −2.00 ± 0.23 | −2.30 ± 0.06 |
| P3 | 72-93 | N-terminal region | −1.50 ± 0.22 | −2.69 ± 0.09 | −2.91 ± 0.07 |
| P4 | 106-126 | N-terminal region | −2.75 ± 0.23 | −3.60 ± 0.13 | −4.25 ± 0.08 |
| P5 | 402-416 | C-terminal region | −0.28 ± 0.18 | −0.45 ± 0.06 | −0.67 ± 0.12 |
| P6 | 417-435 | C-terminal region | −0.62 ± 0.29 | −0.77 ± 0.07 | −0.99 ± 0.29 |
| P7 | 436-455 | C-terminal region | −0.32 ± 0.08 | −0.34 ± 0.13 | −0.48 ± 0.08 |
| P8 | 486-503 | C-terminal region | −0.24 ± 0.07 | −0.38 ± 0.11 | −0.51 ± 0.07 |

ΔD uptake of NOS1AP with **CAPTOR-1** relative to apo (%); negative values indicate protection. Mean ± S.D. of n = 3 independent labeling reactions per condition. Full-length NOS1AP was analyzed apo or after 30 min preincubation with **CAPTOR-1** at a 1:10 protein:compound molar ratio (10 μM:100 μM). Back exchange was corrected with undeuterated and maximally deuterated controls. The eight peptides shown are the statistically significant ligand-responsive peptides, not the complete peptide map.

### Supplementary Table 3. MM/PBSA energy components for NTB24 and CAPTOR-1 bound to the NOS1AP model (kcal/mol)

| Energy term | NTB24 | CAPTOR-1 |
|---|---|---|
| ΔE electrostatic (gas phase) | −31.97 ± 19.54 | −49.61 ± 6.77 |
| ΔE van der Waals (gas phase) | −17.17 ± 3.70 | −28.91 ± 3.24 |
| ΔG polar solvation (PB) | 36.58 ± 16.30 | 58.22 ± 5.65 |
| ΔG nonpolar solvation (surface area) | −2.32 ± 0.44 | −4.06 ± 0.30 |
| **ΔG calculated (sum; entropy not included)** | **−14.87** | **−24.37** |

Calculated with MMPBSA.py from AMBER 20 simulations (ff14SB protein and GAFF2 ligand parameters, TIP3P water), one 100 ns trajectory per system. 41 snapshots were taken from the 80-100 ns interval (every 50th frame). Values are mean ± S.D. across snapshots; snapshots are correlated, so the S.D. is not an independent-replicate error. The calculated ΔG does not include a conformational-entropy term and is not comparable to experimental binding free energies; it is reported only to indicate the direction of change from the parent hit to **CAPTOR-1**.

### Supplementary Table 4. Interface map of the NOS1AP-tau interaction (SPR).

| NOS1AP construct | Full-length 2N4R (1-441) | N-terminal (1-150) | Proline-rich (151-243) | MTBR (244-368) | C-terminal (369-441) |
|---|---|---|---|---|---|
| 1-503 (full-length) | 99.6 ± 5.7 | 4.3 ± 1.3 | 98.5 ± 8.2 | 2.9 ± 2.6 | 2.2 ± 0.5 |
| 1-150 | 97.7 ± 4.9 | 5.1 ± 0.6 | 100.5 ± 2.8 | 2.3 ± 3.2 | 3.9 ± 2.4 |
| 1-126 | 85.1 ± 4.5 | 3.2 ± 3.0 | 89.6 ± 2.2 | 3.5 ± 1.5 | 3.7 ± 1.8 |
| 76-150 | 27.1 ± 5.8 | 3.4 ± 1.8 | 24.1 ± 4.1 | 3.5 ± 1.9 | 3.4 ± 3.2 |
| 1-75 | 3.6 ± 1.9 | 3.6 ± 1.1 | 5.1 ± 1.5 | 3.5 ± 0.8 | 3.3 ± 1.1 |
| 127-503 | 2.8 ± 2.5 | 4.6 ± 2.3 | 4.4 ± 1.1 | 3.6 ± 2.3 | 2.2 ± 1.6 |
| 151-503 | 4.8 ± 2.3 | 2.6 ± 1.3 | 3.8 ± 0.6 | 3.5 ± 1.4 | 1.0 ± 1.2 |

SPR binding response expressed as % of the full-length NOS1AP-full-length tau pair; mean ± S.D. of n = 3 independent preparations. Construct boundaries are residue numbers.

### Supplementary Table 5. Phosphoproteomics of PFF-exposed neurons (AD-model): CAPTOR-1, NOS1AP knockdown and resistant NOS1AP

| Phosphosite | Family | Control | AD | AD + CAPTOR-1 | AD + NOS1AP CRISPRi | AD + CRISPRi + resistant NOS1AP + CAPTOR-1 |
|---|---|---|---|---|---|---|
| Tau pT217 | Primary (tau) | +0.08 ± 0.14 | +1.37 ± 0.21 | +0.54 ± 0.20 | +0.49 ± 0.18 | +1.22 ± 0.16 |
| Tau pT231 | Primary (tau) | +0.02 ± 0.13 | +1.12 ± 0.18 | +0.40 ± 0.19 | +0.38 ± 0.22 | +1.02 ± 0.15 |
| Tau pS202 | Primary (tau) | −0.01 ± 0.14 | +1.03 ± 0.12 | +0.45 ± 0.18 | +0.28 ± 0.20 | +0.84 ± 0.12 |
| GSK3β pY216 | Exploratory | +0.01 ± 0.23 | +0.80 ± 0.19 | +0.26 ± 0.09 | +0.24 ± 0.20 | +0.58 ± 0.13 |
| p38α pT180/Y182 | Exploratory | +0.09 ± 0.18 | +0.85 ± 0.14 | +0.32 ± 0.11 | +0.21 ± 0.18 | +0.81 ± 0.17 |
| CaMKIIα pT286 | Exploratory | +0.01 ± 0.13 | −0.69 ± 0.15 | −0.27 ± 0.13 | −0.06 ± 0.19 | −0.49 ± 0.21 |
| CREB pS133 | Exploratory | −0.05 ± 0.17 | −0.65 ± 0.16 | −0.20 ± 0.13 | −0.28 ± 0.17 | −0.61 ± 0.21 |
| SYN1 pS9 | Exploratory | +0.02 ± 0.20 | −0.59 ± 0.20 | −0.27 ± 0.11 | −0.08 ± 0.12 | −0.31 ± 0.16 |
| PSD-95 pS73 | Exploratory | −0.03 ± 0.15 | −0.35 ± 0.24 | −0.13 ± 0.16 | −0.14 ± 0.15 | −0.38 ± 0.11 |
| NOS1 pS1417 | Exploratory | −0.01 ± 0.23 | +0.77 ± 0.16 | +0.26 ± 0.21 | +0.20 ± 0.12 | +0.54 ± 0.29 |
| FYN pY420 | Exploratory | −0.08 ± 0.13 | +0.53 ± 0.11 | +0.27 ± 0.15 | +0.17 ± 0.17 | +0.53 ± 0.16 |
| GluN2B pY1472 | Exploratory | −0.01 ± 0.13 | +0.51 ± 0.20 | +0.35 ± 0.14 | +0.16 ± 0.11 | +0.55 ± 0.10 |

**P value, Holm-adjusted within family (unadjusted P in parentheses)**

| Phosphosite | Family | AD versus control | AD + CAPTOR-1 versus AD | AD + CRISPRi versus AD | Resistant + CAPTOR-1 versus AD + CAPTOR-1 |
|---|---|---|---|---|---|
| Tau pT217 | Primary (tau) | 0.0022 (0.0022) | 0.0059 (0.0039) | 0.0070 (0.0035) | 0.0003 (< 0.0001) |
| Tau pT231 | Primary (tau) | 0.0005 (0.0002) | 0.0017 (0.0006) | 0.0070 (0.0042) | 0.0007 (0.0003) |
| Tau pS202 | Primary (tau) | 0.0004 (0.0001) | 0.0059 (0.0029) | 0.0030 (0.0010) | 0.0079 (0.0079) |
| GSK3β pY216 | Exploratory | 0.064 (0.021) | 0.068 (0.0096) | 0.088 (0.027) | 0.013 (0.0018) |
| p38α pT180/Y182 | Exploratory | 0.043 (0.0086) | 0.068 (0.0085) | 0.050 (0.0062) | 0.0063 (0.0007) |
| CaMKIIα pT286 | Exploratory | 0.010 (0.0011) | 0.072 (0.014) | 0.077 (0.015) | 0.19 (0.075) |
| CREB pS133 | Exploratory | 0.032 (0.0040) | 0.025 (0.0027) | 0.071 (0.012) | 0.0087 (0.0011) |
| SYN1 pS9 | Exploratory | 0.048 (0.012) | 0.082 (0.027) | 0.068 (0.0097) | 0.57 (0.57) |
| PSD-95 pS73 | Exploratory | 0.064 (0.028) | 0.085 (0.043) | 0.088 (0.062) | 0.072 (0.012) |
| NOS1 pS1417 | Exploratory | 0.040 (0.0057) | 0.068 (0.0093) | 0.018 (0.0020) | 0.17 (0.042) |
| FYN pY420 | Exploratory | 0.040 (0.0062) | 0.072 (0.015) | 0.088 (0.022) | 0.13 (0.026) |
| GluN2B pY1472 | Exploratory | 0.064 (0.030) | 0.24 (0.24) | 0.088 (0.023) | 0.19 (0.063) |

Paired t-tests on donor means (n = 4 donors). Holm adjustment is applied within each contrast across the three tau sites (primary family) or across the nine signaling nodes (exploratory family). "Resistant" = NOS1AP CRISPRi neurons re-expressing the drug-resistant NOS1AP variant (Y16A/F42A) and treated with **CAPTOR-1**.

### Supplementary Table 6. In vitro ADME properties of CAPTOR-1

| Property | Value | Unit |
|---|---|---|
| Kinetic solubility, pH 7.4 | 86.0 ± 9.0 | uM |
| Mouse liver microsome t1/2 | 54.0 ± 6.0 | min |
| Human liver microsome t1/2 | 68.0 ± 8.0 | min |
| PAMPA-BBB permeability (Pe) | 8.1 ± 0.9 | $10^{-6}$ cm/s |
| hERG inhibition | 18.0 ± 4.0 | % inhibition at 10 uM |

Mean ± S.D. Assay conditions and n per assay.
